# Unveiling the Power of High-Dimensional Cytometry Data with cy*CONDOR*

**DOI:** 10.1101/2024.02.29.582727

**Authors:** Charlotte Kroeger, Sophie Müller, Jacqueline Leidner, Theresa Kröber, Stefanie Warnat-Herresthal, Jannis Bastian Spintge, Timo Zajac, Aleksej Frolov, Caterina Carraro, Simone Puccio, Joachim L Schultze, Tal Pecht, Marc D Beyer, Lorenzo Bonaguro

## Abstract

High-dimensional cytometry (HDC) is a powerful technology for studying single-cell phenotypes in complex biological systems. Although technological developments and affordability have made HDC broadly available in recent years, technological advances were not coupled with an adequate development of analytical methods that can take full advantage of the complex data generated. While several analytical platforms and bioinformatics tools have become available for the analysis of HDC data, these are either web-hosted with limited scalability or designed for expert computational biologists, making their use unapproachable for wet lab scientists. Additionally, end-to-end HDC data analysis is further hampered due to missing unified analytical ecosystems, requiring researchers to navigate multiple platforms and software packages to complete the analysis.

To bridge this data analysis gap in HDC we developed *cyCONDOR*, an *easy-to-use* computational framework covering not only all essential steps of cytometry data analysis but also including an array of downstream functions and tools to expand the biological interpretation of the data. The comprehensive suite of features of *cyCONDOR*, including guided pre-processing, clustering, dimensionality reduction, and machine learning algorithms, facilitates the seamless integration of *cyCONDOR* into clinically relevant settings, where scalability and disease classification are paramount for the widespread adoption of HDC in clinical practice. Additionally, the advanced analytical features of *cyCONDOR*, such as pseudotime analysis and batch integration, provide researchers with the tools to extract deeper insights from their data. We used *cyCONDOR* on a variety of data from different tissues and technologies demonstrating its versatility to assist the analysis of high dimensionality data from preprocessing to biological interpretation.

## Introduction

The rapid development of high-dimensionality cytometry (HDC) methods has revolutionized how we can analyze millions of cells from thousands of complex tissues. Mainly driven by immunological research, where the heterogeneity of cell types and the growing number of cell states particularly benefits from these high-dimensionality techniques ^1^, HDC is now extremely robust and routinely employed to measure simultaneously up to 50 markers at single-cell resolution, making it instrumental not only in immunological research, but increasingly in other disciplines such as microbiology, virology, or neurobiology ^2^. The main technologies employed in this field are high-dimensionality flow cytometry (HDFC) ^3^, total spectrum flow cytometry (SpectralFlow) ^4^, Cytometry by time of flight or mass cytometry (CyTOF) ^5^ and proteogenomics (CITE-seq/Ab-seq) ^6^. These antibody-based methods allow not only the detection of intra- and extra-cellular proteins but also the specific identification of post-translational modifications, adding an important functional layer to nucleotide-based methods (e.g. single-cell RNA sequencing). Particularly the cytometry-based methods are characterized by significant throughput allowing the measurement of millions of cells per sample ^1^.

While HDCs come with many advantages and opportunities, their high-dimensionality also comes with challenges, of which a major one is the application of conventional analytical approaches that rely on consecutive gating based on one or two parameters at a time. It has been shown recently that conventional analytics are prone to miss the intricate relationships and patterns that exist within high-dimensional datasets, which can lead to incomplete and potentially misleading interpretations ^1^. Effectively harnessing the full potential of HDC datasets requires an unbiased perspective and the ability to operate without the need for prior knowledge ^1^. Along these lines specialized bioinformatics tools were developed capable of navigating the complexity of HDC datasets and extracting meaningful insights without relying on pre-existing assumptions.

In the last few years, several approaches besides commercial software have provided the cytometry community with tools to investigate HDC data using data-driven approaches commonly used by the single-cell transcriptomics community. *Cytofkit* ^7^, a pioneering project that ceased development in 2017, *SPECTRE* ^8^ and *Catalyst* ^9^ have extensively contributed to the current standards of HDC data analysis. Nevertheless, these tools do not yet provide an end-to-end ecosystem for HDC data analysis. Complementary, several non-academic projects, such as *Cytobanks* or *Cytolytics* provide feature-rich tools, often with an intuitive graphical user interface (GUI) for the guided analysis of HDC data. These implementations, while extremely useful for wet-lab scientists, often fail to scale well with large datasets.

We hypothesized that an integrated, simple to use, end-to-end ecosystem for HDC data analysis would overcome current shortcomings and enable HDC users to take full advantage of the high dimensionality of the data. The solution is an integrated ecosystem (1) unifying different algorithms for a diverse set of analyses under a united data structure; (2) being able to analyze a high number of cells/samples optimized for consumer hardware but deployable on high-performance computers (HPCs); and (3) designed with a focus on data interpretation and visualization.

Here we present *cyCONDOR* (<u>github.com/lorenzobonaguro/cyCONDOR</u>) for the analysis of HDC data. Our tool provides an integrated ecosystem for the analysis of CyTOF, HDFC, SpectralFlow and CITE-seq data in R in a unified format designed for its ease of use by non-computational biologists (**Figure 1a**). *cyCONDOR* offers a comprehensive data analysis toolkit encompassing data ingestion and transformation, batch correction, dimensionality reduction, and clustering, along with streamlined functions for data visualization, biological comparison, and statistical testing. Its advanced features include deep learning algorithms for automated annotation of new datasets and classification of new samples based on clinical characteristics (**Figure 1b**). Additionally, *cyCONDOR* can infer the pseudotime of continuous biological processes to investigate developmental states or disease trajectories ^10^ (**Figure 1b**). Compared to other currently available toolkits, *cyCONDOR* provides the most comprehensive collection of analysis algorithms and the most interpretable data format (**Figure S1a**). Furthermore, the entire *cyCONDOR* ecosystem was designed to be scalable to millions of cells while being still usable on common hardware (**Figure S1b**). We used *cyCONDOR* on a variety of private and public datasets showing seamless compatibility with all tested cytometry data formats. We made *cyCONDOR* available in R as a standalone package or as containerized environments easily deployed on local hardware or HPCs. With *cyCONDOR*, we provide an ecosystem that allows the end user to fully exploit the potential of HDC methods.

**Figure 1:**
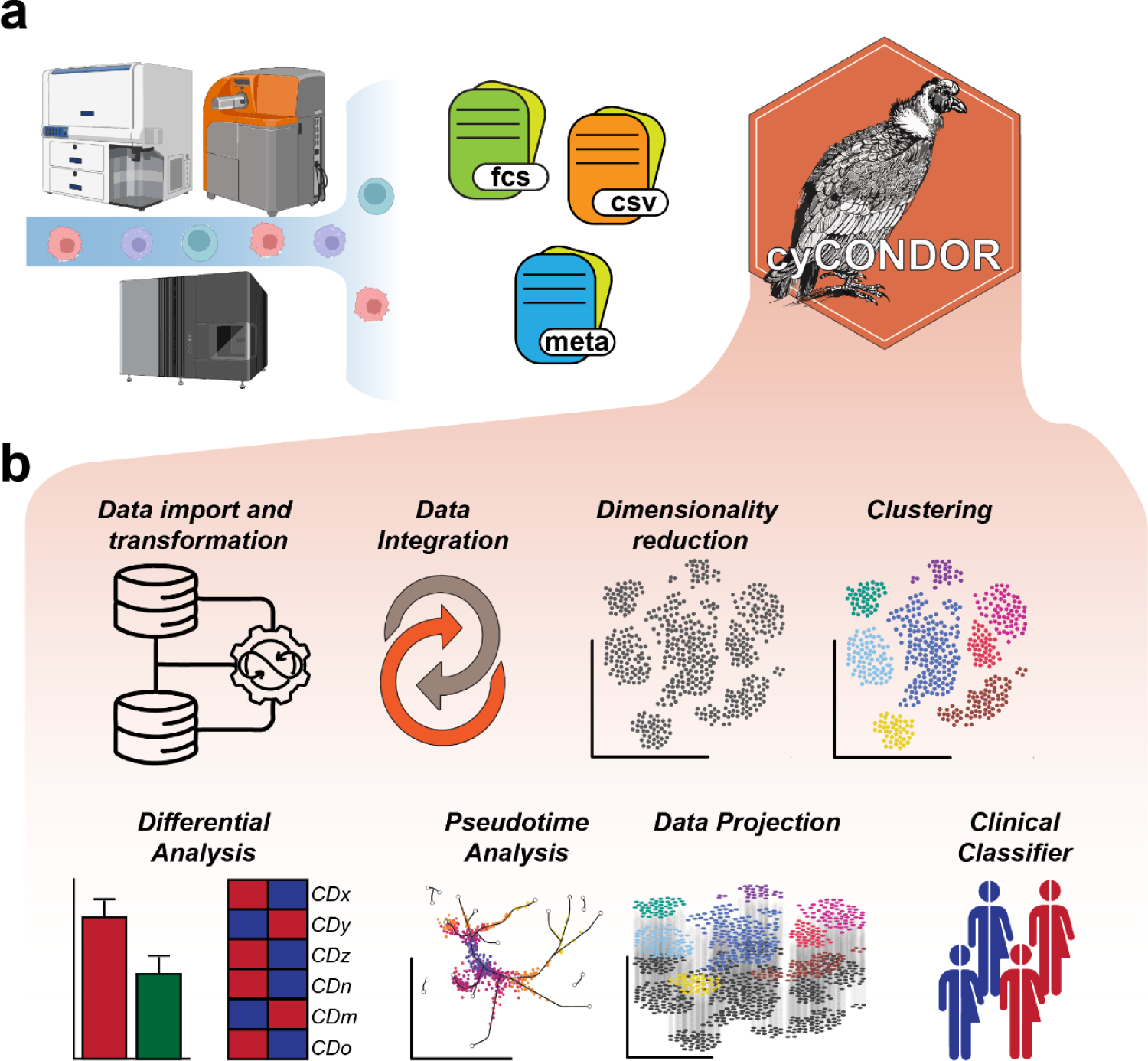
Overview of the *cyCONDOR* ecosystem. **a,** The *cyCONDOR* ecosystem accepts HDC data from a variety of technologies combined with sample annotation**. b,** The ecosystem covers a broad variety of analytical tasks, from data import and transformation to ML-based sample classifiers.

## Results

### *cyCONDOR* provides a versatile workflow for data pre-processing

cyCONDOR offers a suite of microservices for data import and pre-processing to make use of a versatile set of data input formats in HDC (**Fig. 1a**) and to provide the necessary data pre-processing prior to an integrated higher-level data analysis (**Fig 1b**). As default input data format for the *cyCONDOR* workflow, either Flow Cytometry Standard files (.fcs) or Comma-separated values files (.csv) are used, which can be exported by current acquisition or flow cytometry data analysis software such as FlowJo (www.flowjo.com, **Supplementary Information**). In addition, metadata describing the dataset are also imported. Users may choose to include all recorded events in the output files or apply upfront broad gating to reduce dataset size. We recommend applying basic gating prior to *cyCONDOR* to exclude debris and doublets, thereby minimizing the downstream computational demand. This simple pre-filtering step removes irrelevant events and significantly reduces computational requirements, enabling the analysis of even large datasets on consumer-grade hardware.

Following data import, *cyCONDOR* provides a comprehensive end-to-end ecosystem of HDC data pre-processing and analysis (**Figure 2a, S2a**). In the following sections, we will exemplify the use of *cyCONDOR* for the analysis of HDC data. All output shown here is the result of built-in functions and can be generated for any other dataset with minimum bioinformatics knowledge. In the following example, we explore a human PBMCs dataset ^11^ to exemplify the first steps of a *cyCONDOR* analysis. This dataset, including 27 protein markers, provides a broad phenotyping of the main circulating immune cells in human peripheral blood derived from people living with HIV (PLHIV, Dis) and uninfected individuals (controls, Ctrl). *cyCONDOR* exploratory data analysis starts with data loading and transformation to ensure a distribution of values compatible with downstream investigations (<u>see Methods for details</u>) (**Figure 2a, S2a**). To initially visualize the underlying data structure and to explore whether the distribution of samples is linked to factors like biological group, age, sex or time of sampling, principal component analysis (PCA) is performed on pseudobulk samples calculated as the sum of protein expression of all cells (<u>details in Methods</u>, **Figure 2b**). The average expression for each marker on a sample level can be inspected to help identifying the main drivers of the observed biological differences for example between two defined groups within the dataset (**Figure 2c**). In our example, we see a general decrease in T cell markers (e.g. CD3 and CD4) in PLHIV versus Ctrl and an overall increased expression of monocytes markers (e.g. CD14 and HLA-DR), which can be interpreted as either an increased expression of those markers in PLHIV cells or, most likely as a shift in the relative frequency of cells in HIV patients (**Figure 2c**). When analyzed at the single-cell level (**Figure S2b**), the dataset reveals patterns that can be further elucidated by visualizing the loadings of the most relevant principal components (**Figure S2c**) which - in our example - revealed T cell-associated markers CD27, CD3, CD127 and CD8. Further, to reduce the dimensionality of the dataset to a bi-dimensional space, *cyCONDOR* provides the implementation of two non-linear dimensionality reduction algorithms, Uniform Manifold Approximation and Projection (UMAP ^12,13^) and t-distributed Stochastic Neighbor Embedding (tSNE ^14^) as they both have different advantages (<u>see methods for details</u>). UMAP ^12^ dimensionality reduction can be performed (**Figure 2d**), and visualized as a two-dimensional scatter plot, colored for any variable of interest (e.g. experimental group or date, **Figure 2d**) or visualized as a density plot, to highlight the distribution of the cells in the latent space (**Figure s2d**). The two-dimensional UMAP embedding can also be used to visualize the expression of the individual protein markers (**Figure S2e**). Additionally, for unsupervised non-linear dimensionality reduction tSNE is implemented in *cyCONDOR* (**Figure S3e**).

**Figure 2:**
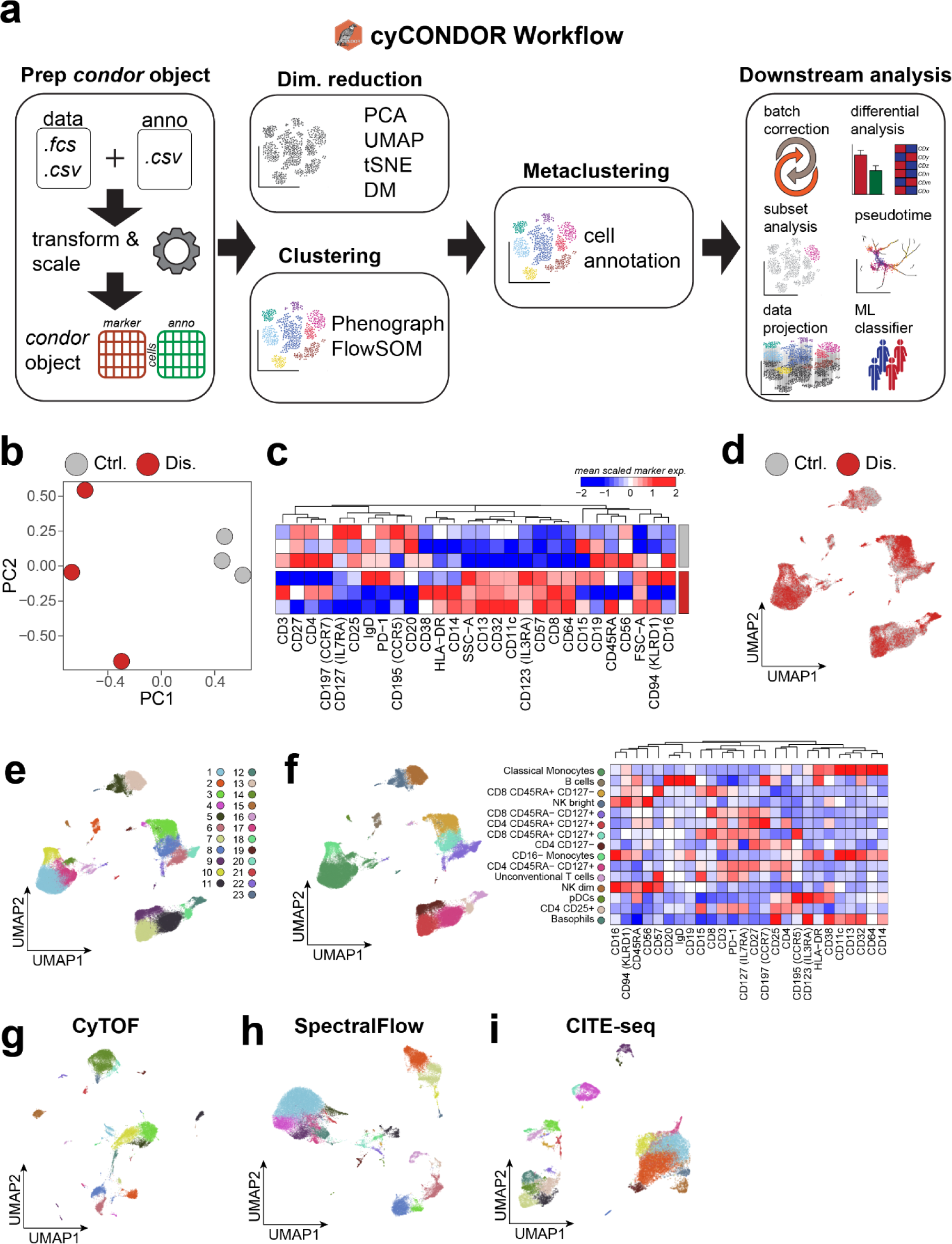
*cyCONDOR* workflow for data pre-processing and annotation. **a,** Schematic view of the first steps of *cyCONDOR* analysis, from data ingestion to cell labelling. **b,** Pseudobulk Principal Component Analysis (PCA) colored by experimental groups. **c,** Heatmap showing mean marker expression for each samples, column order is defined by hierarchical clustering. **d,** UMAP colored by experimental group. **e,** UMAP colored according to Phenograph clustering. **f,** UMAP colored according to cell type annotation and heatmap of mean marker expression for each cell type. **g,** UMAP visualization of SpectralFlow data colored by Phenograph clustering. **h,** UMAP visualization of CyTOF data colored by Phenograph clustering. **i,** UMAP visualization of CITE-seq data colored by Phenograph clustering.

To assign cell type labels *cyCONDOR* provides two different clustering algorithms Phenograph ^15^ and FlowSOM ^16^ integrated here into the cyCONDOR workflow providing different data output formats (**Figure 2e, S3a-d**). The combination of FlowSOM for fast knowledge-based clustering (**Figure S3b-d**) and Phenograph (**Figure 2e, S3a**) enables data-driven identification of major cell lineages and the potential discovery of novel cell states through slower but fine-grained clustering ^17^. To ease the biological annotation of the clusters *cyCONDOR* provides an automated heatmap visualization of the average gene expression of each cluster (**Figure S3a, S3d**). As a next step, users can manually label each cluster according to prior knowledge in the field concerning identity (**Figure 2f**). Annotated clusters and embeddings are the starting point for further downstream analysis provided within *cyCONDOR*. To illustrate the applicability of the *cyCONDOR* ecosystem not only to HDFC data (exemplified so far in Figure 2) we performed data transformation, dimensionality reduction and clustering also on published CyTOF (**Figure 2g, S3f, S3g**), Spectral Flow and (**Figure 2h, S3h, S3i**) CITE-seq datasets (**Figure 2i, S3j, S3k**) showing general applicability of *cyCONDOR* to all major cytometry data types.

### *cyCONDOR* provides correction of technical variance across projects, time, datasets, instruments, or sites

Similarly to other high dimensionality techniques (e.g. RNA sequencing or proteomics), HDC methods suffer from the presence of technical variation making it challenging to integrate datasets generated from different projects, datasets, instruments, sites or at different times ^18^. When compared to other high-dimensional methodologies, HDC falls behind, since the parameter space is increasingly inflated with new technical opportunities, literally allowing any combination of antibody and detection reagents such as fluorochromes in flow cytometry in addition to increasing opportunities for diverse configurations of instruments and instrument performances ^18^. To cope with these developments, we implemented *Harmony* ^19^ in *cyCONDOR* for batch alignment over multiple sources of technical variation. *Harmony* was introduced as a tool for correction of technical variation in single-cell RNA sequencing data ^20^ but its applicability can be easily generalized to other single-cell methods such as HDC with the only requirement of a normal distribution of the parameters to be harmonized (e.g. normalized fluorescence intensity or principal components).

*cyCONDOR* offers the option to apply *Harmony* variance correction on protein expression or principal components (**Figure 3a, S4a**). Although the direct harmonization of fluorescence intensities can provide important information on the source of variability, corrected intensities should be used carefully, especially in the analysis of differential expression across experimental groups ^21^.

**Figure 3:**
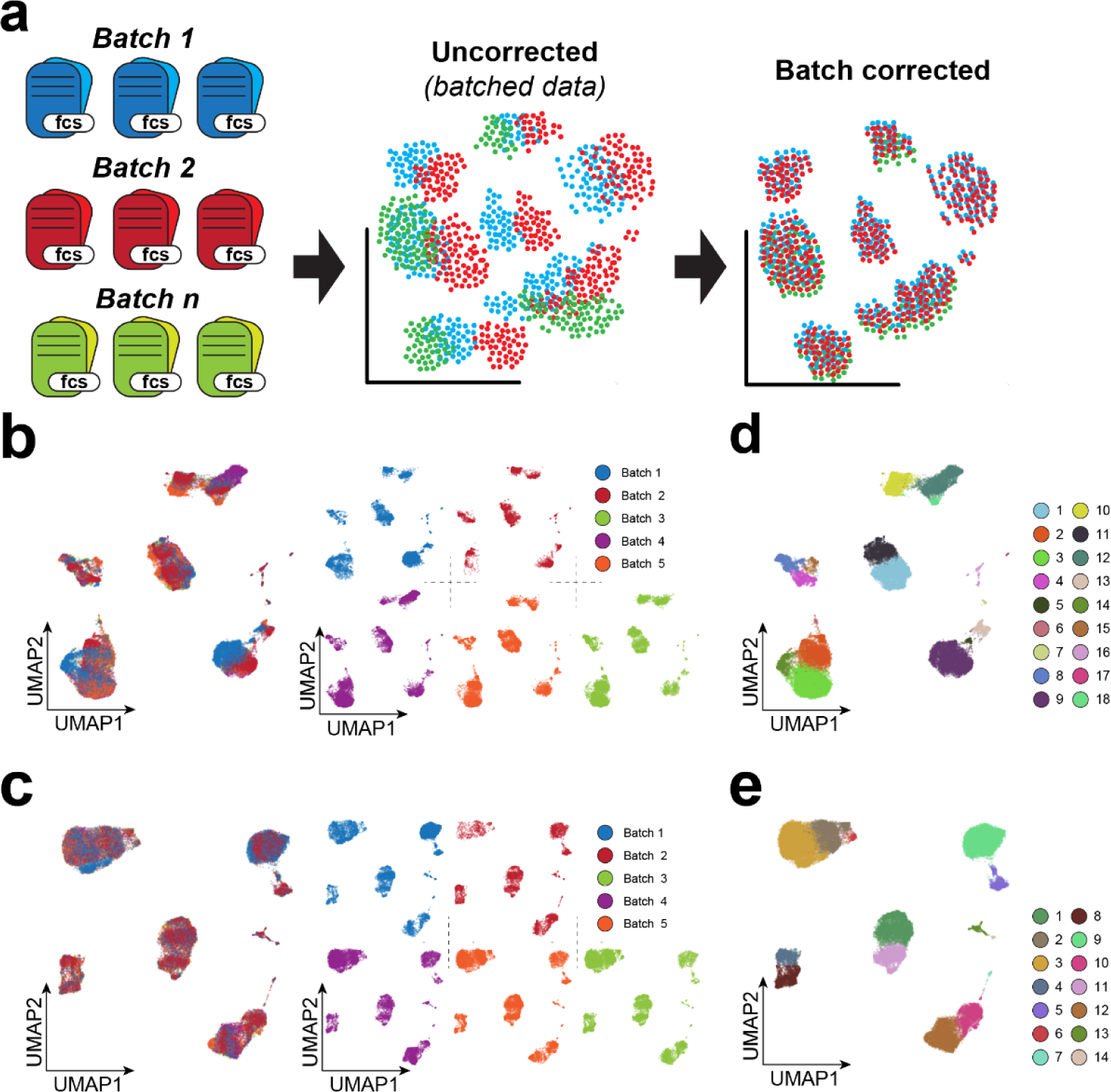
Technical differences between batches can be mitigated with *cyCONDOR*. **a,** Schematic overview of the batch correction workflow implemented in *cyCONDOR*. **b,** Original UMAP colored according to the experimental batch (left) and split by the experimental batch (right). **c,** Batch corrected UMAP colored according to the experimental batch (left) and split by the experimental batch (right). **d,** Original UMAP colored by Phenograph clustering. **e,** Batch corrected UMAP colored by Phenograph clustering.

Here, we showcase the performance of technical variation correction provided by *cyCONDOR* on a 27-color flow cytometry dataset where healthy controls were measured at five different time points across three months with adjustments on the instrument settings due to inconsistencies in instrument performance (*unpublished data*). Such example showcases a rather common situation in clinical studies where patient samples are processed over several weeks or months if not years. Instruments performance quality control (QC) and automatic adjustments ^22,23^ can help to reduce those biases but in high dimensionality data, those are difficult to be fully resolved. This can be illustrated by representing the data in a UMAP, a non-linear dimensionality reduction, which reveals a high degree of separation between different experimental dates (**Figure 3b**), exemplified also by a low Local Inverse Simpson’s Index (LISI) score ^19^ (**Figure S4b**). *Harmony* correction on all calculated principal components mitigates the technical variance in the UMAP embedding showing a more homogeneous distribution of each batch in the clusters. (**Figure 3c**). This improvement was quantified by calculating the LISI score showing a remarkable increase compared to pre-correction scores (**Figure S4b**)

To further investigate the batch effect across dates, Phenograph clustering was performed on both non-corrected PCs (**Figure 3d**) and *Harmony*-corrected PCs (**Figure 3e**) with identical resolution settings. Clustering based on not-corrected principal components (PCs) leads to the identification of 18 clusters, but further inspection revealed that most of them are date-specific - most prominently cluster 6, 14, 15, 18 (**Figure S4c**). After *Harmony* batch correction, only cluster 6 appears to be still specific for batch three (**Figure S4d**). Investigating this persisting difference between batches at the level of individual samples revealed that the majority of the cells in cluster 6 derive from one sample (belonging to *batch 3,* **Figure S4e**), showing our approach was successful in removing unwanted technical variability while preserving the biological difference between samples.

### Pseudotime Projection-Based Trajectory Inference allows the dissection of developmental programs

A valuable insight enabled by single-cell level analysis over bulk analysis is the capacity to investigate continuous developmental trajectories in complex tissues ^10^. While HDC provides sufficient resolution for this type of analysis, conventional analysis approaches based on classical gating of the data can only capture discrete cell states but fail to capture the whole scope of continuous processes ^24^. The technical and conceptual framework of *cyCONDOR* allows to integrate approaches which are defining pseudotimes as a proxy for continuous developmental trajectories based for example on cluster-based minimum spanning trees as they have been realized by the *slingshot* algorithm ^25^ to predict pseudotime in single-cell data. This addition to *cyCONDOR* opens the potential to investigate complex transitional states in HDC data.

To illustrate the potential of pseudotime analysis on HDC data we analyzed a bone marrow CyTOF dataset from Bendal and colleagues ^26^ with a dimensionality of 32 protein markers to visualize the developmental trajectories of hematopoietic stem cells (HSCs) to monocytes and plasmacytoid dendritic cells (pDCs).

The first step of this analysis includes the annotation of the dataset (as described in **Figure 2**) and the subsetting for the myeloid lineage (**Figure 4a, S5a**). The subsetting function is especially useful for a high-resolution analysis of highly heterogeneous tissues, such as the bone marrow. Bone marrow data was pre-processed and each Phenograph cluster was annotated according to the expression of hallmark proteins (**Figure 4b, S5b, S5c)**. From the entire cellular space we focused on the myeloid cell compartment including monocytes and plasmacytoid dendritic cells (pDCs) (**Figure 4c**) to define their differentiation trajectories. Dimensionality reduction and clustering were reiterated on the selected cell compartment to increase the resolution of cell types and states, resulting in 15 clusters (**Figure S5d**). Importantly, the subset data was not re-scaled for clustering and dimensionality reduction (as it is e.g. performed in standard single-cell transcriptomics workflow such as Seurat ^27^ or Scanpy ^28^) to avoid any overrepresentation of proteins not expressed. Finally, each cluster was labeled according to the expression of lineage proteins (**Figure 4c, S5e**) revealing the presence of a common myeloid progenitor (CMPs) cluster which was not resolved before subsetting.

**Figure 4:**
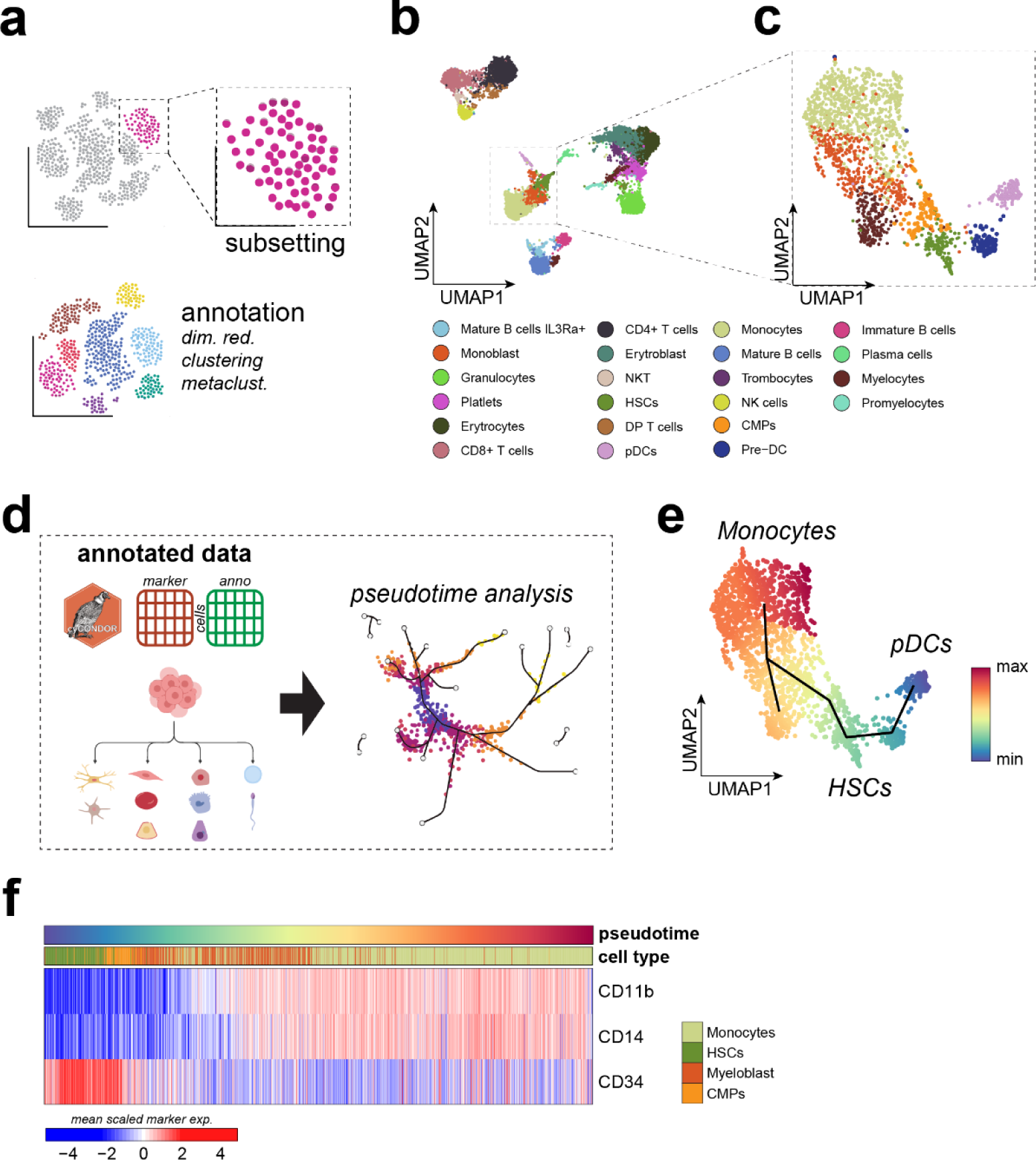
Pseudotime inference on cytometry data helps to describe continuous developmental processes. **a,** Schematic overview of the subsetting workflow implemented in *cyCONDOR*. **b,** UMAP of all BM cells colored by annotated cell type. **c,** UMAP of the subset of monocytes, pDCs and their progenitors colored by annotated cell type. **d,** Schematic overview of the pseudotime inference workflow implemented in *cyCONDOR*. **e,** UMAP colored according to the inferred pseudotime of the predicted trajectories. **f,** Heatmap of protein expression in cells belonging to the monocytes trajectory ordered according to the inferred pseudotime.

Within the *cyCONDOR* ecosystem, we can infer pseudotime and trajectories on the filtered dataset using the PCs or UMAP coordinates as an input (**Figure 4d, S6a**). In the slingshot function, it is possible to force the pseudotime to start and end at specific clusters. However, we suggest allowing slingshot to infer the best starting and ending point of the trajectory and corroborate the results with domain knowledge for the analysis ^25^. In our example, *slingshot* unbiasedly predicted a developmental trajectory starting at one of the pDCs clusters via the HSC cluster towards the monocyte clusters, where it branched at the level of myeloblasts (**Figure 4e**). Incorporating prior biological knowledge, namely that HSCs are at the starting point of cell differentiation within the myeloid compartment, the interpretation of the pseudotime analysis would suggest that pDC development trajectory is distinct from monocyte development and that the different monocyte subsets share a common differentiation path from HSCs to myeloblasts and subsequently into monocytes (**Figure 4f, S6b**). In the first branch, leading from HSCs to monocytes, we observed a gradual decline of HSCs markers (e.g. *CD34*) and an increased expression of monocyte markers such as *CD11b* and *CD14* (**Figure 4f**). In contrast, the developmental trajectory from HSCs to pDCs was defined by a decline of *CD34* and *HLA-DR* expression and an increased expression of *CD123*, a hallmark protein for pDCs (**Figure S6b**). This CyTOF dataset exemplifies the value of pseudotime analysis of HDC data beyond sequencing-based single cell technologies, allowing a more fine-granular analysis of cellular differentiation states for example in the hematopoietic system, the immune system, but potentially also in cancer or other renewing tissues.

### *cyCONDOR* empowers visual and statistical comparison between experimental groups

Many HDC analyses aim to investigate the biological difference between two or more experimental groups or conditions. Despite the availability of tools for pre-processing HDC data ^7–9,29^, comprehensive frameworks for in-depth visualization and statistical testing to compare multiple biological groups remain limited. With *cyCONDOR* we provide a set of easy-to-use functions to compare cell frequencies and protein expression across multiple experimental groups (**Figure 5a, S7a**).

**Figure 5:**
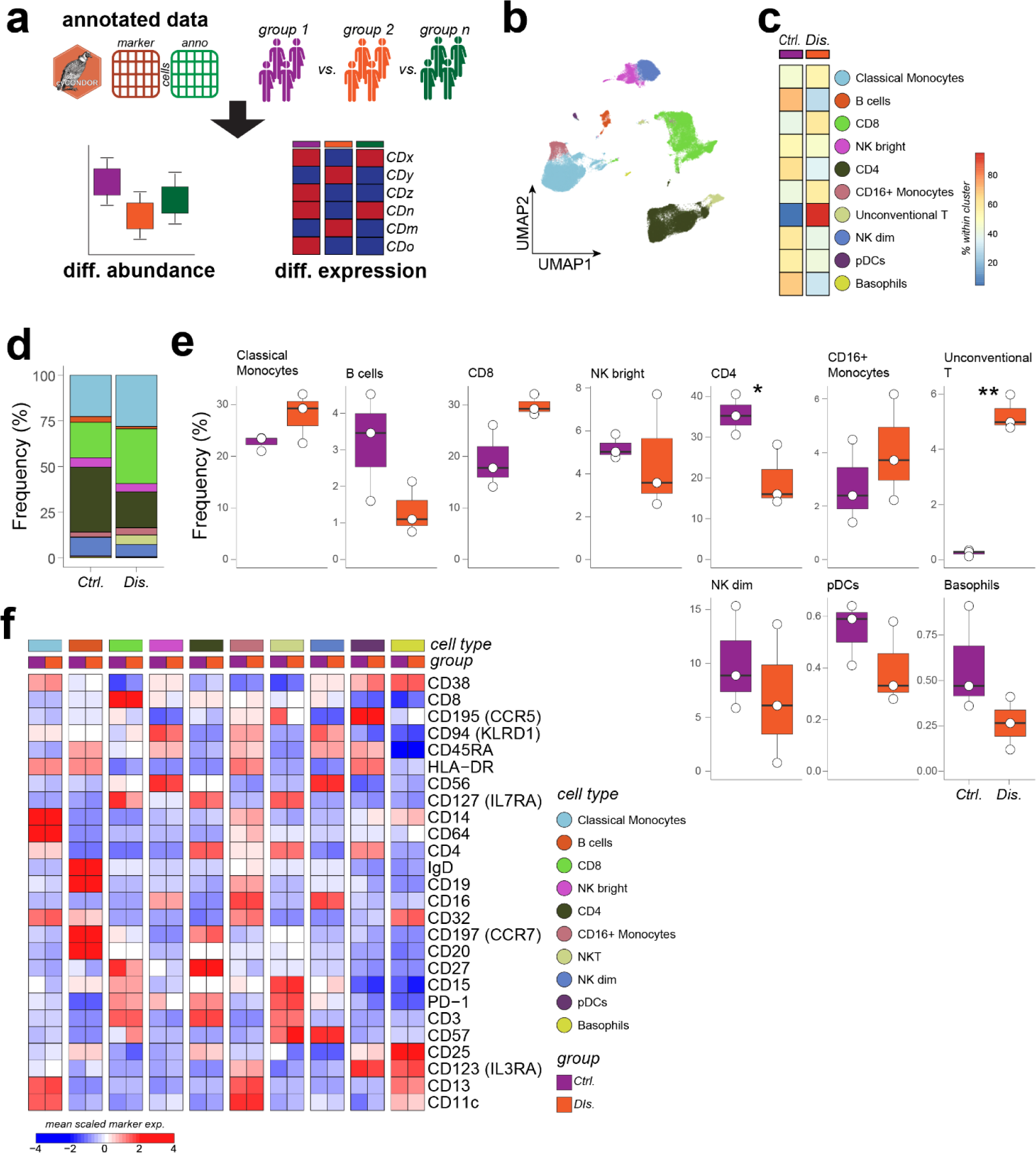
*cyCONDOR* provides accessible function for differential analysis. **a,** Schematic overview of the differential analysis workflow. **b,** UMAP of the PBMCs dataset colored by annotated cells type. **c,** Confusion matrix of the annotated cell types split by experimental group. **d,** Stacked barplot of the cellular frequencies of the annotated cell types split by experimental groups. **e,** Boxplot of the frequency of each annotated cell type split by experimental group. **f,** Heatmap of the average expression of each marker split by cell type and experimental group. Statistical significance was calculated with a t-test with default settings, * p < 0.05, ** p < 0.01, *** p < 0.001.

To exemplify these built-in features of *cyCONDOR* we re-analyzed a subset of our previously published dataset on chronic HIV ^11^. Pre-processing of the dataset, including data transformation, dimensionality reduction, clustering and cell annotation (as described in **Figure 2**) revealed the presence of the expected cell populations in PBMCs (**Figure 5b**). At a glance, the contribution of each experimental group to each cell type (**Figure 5c**) or cluster (**Figure S7b, S7c**) can be visualized as confusion matrix. *cyCONDOR* provides stacked bar plots as a second integrated visualization approach to compare cell compositions per group (**Figure 5d, S7d**). Interestingly, already at this level a reduced frequency of B cells and CD4+ T cells and an increased frequency of monocyte and non-conventional T cells was observed, as expected in individuals with chronic HIV infection (**Figure 5c, 5d,** ^11^). Already these simple visualization approaches provide fast and easily interpretable overviews. Yet, they do not address potential sample outliers or provide statistical testing.

Cell frequencies at the sample level separated by groups are visualized with a built-in *cyCONDOR* function generating boxplots for each cell type or cluster for each sample group individually (**Figure 5e, S5e**), providing tabular output with summary statistics and several options for statistical testing (see methods for details).

Differential protein expression between conditions of interest can also be investigated with a built-in function of *cyCONDOR* by providing only the cell labels to be used for the categorization (e.g. clustering or cell types) and the biological grouping. The result is visualized as a heatmap of the average gene expression across groups and cell types, showing for example a decreased expression of the naive T cell markers CD127 and CD27 and an increased expression of the senescence marker CD57 in CD8+ T cells of PLHIV (**Figure 5f, S8a**).

Overall *cyCONDOR* provides a diverse collection of easy-to-use functions to investigate the biological differences between experimental groups to cover a wide-range of statistical comparisons and visualization needs.

### Continuous learning and scalability in HDC leveraging data projection with cyCONDOR

Considering the high scalability and the continuously increasing affordability of HDC, it is of utmost importance to establish an analytical pipeline designed to be scalable to the analysis of thousands of samples and millions of cells. Given the widespread adoption of HDC as the primary readout for numerous longitudinal population-wide or clinical studies, a real-time processing of the growing datasets upon each novel data acquisition is impractical and inefficient. With *cyCONDOR* we propose a two-step approach for continuous learning from new data (**Figure 6a, S9a**). As a first step, a representative set of samples will be used to generate the initial cell state and protein expression model (**Figure 6b, S9b**). This initial model should be as representative as possible for the variability of the samples and their cell populations to be analyzed and the specific scientific question to be answered ^30^. As a second step with a transfer-learning approach, newly generated data will be projected onto the annotated reference for an efficient cell annotation of new data.

**Figure 6:**
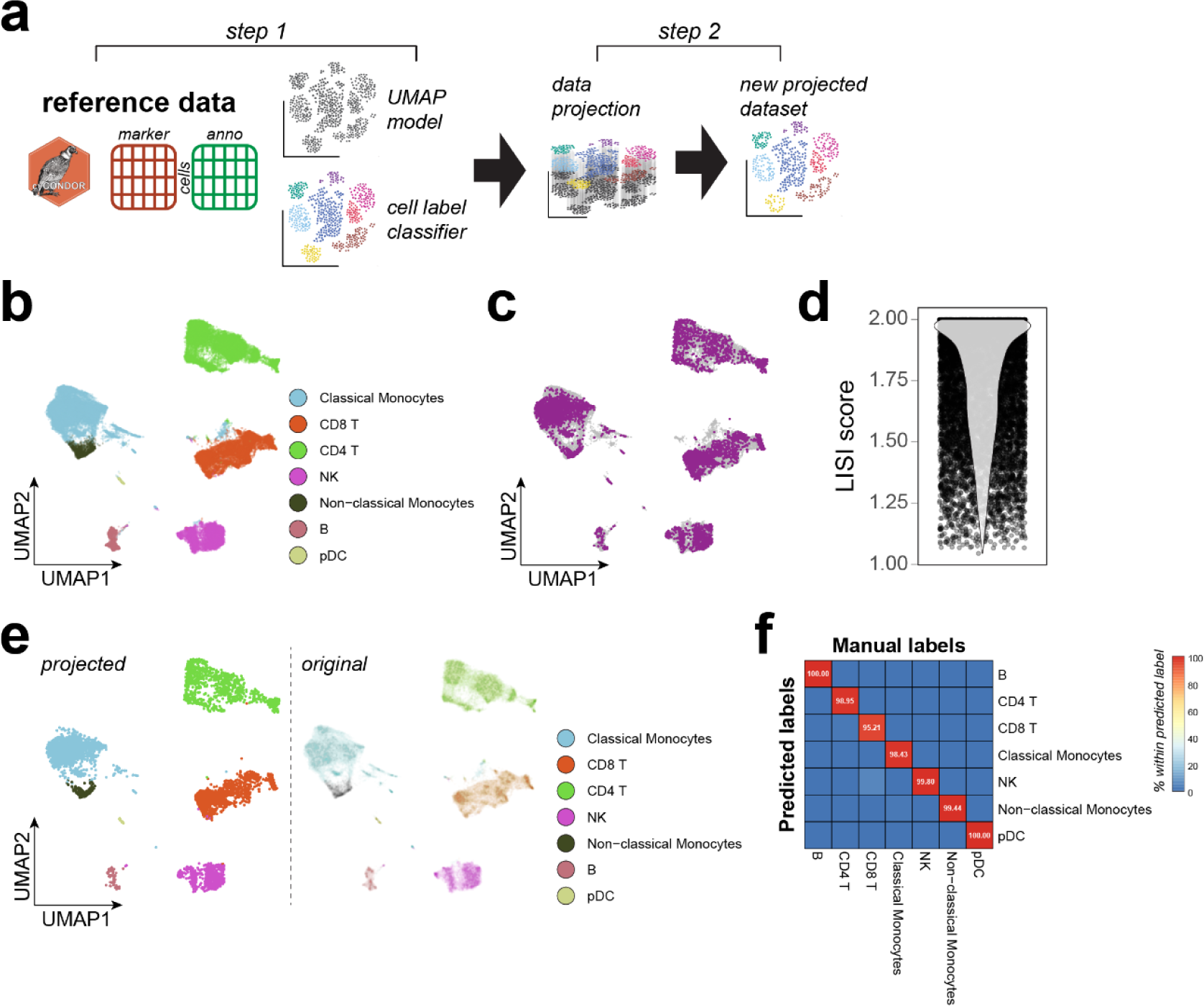
Batch alignment allows accurate analysis of longitudinal data. **a,** Schematic overview of the data projection workflow implemented in *cyCONDOR*. **b,** UMAP visualization of the training dataset colored according to the annotated cell type. **c,** UMAP overlapping the projected data (purple) to the training dataset (grey). **d,** LISI scores calculated between training and projected data. **e,** Left: UMAP visualization of the projected data colored according to the predicted cell types, right: UMAP of the original data colored by cell label used to train UMAP model and kNN classifier. **f,** confusion matrix comparing the manual annotation of the projected data with the predicted cell labels.

Following the principles described above (**Figure 2**), a representative set of samples is processed by dimensionality reduction clustering and cluster annotation. Next, the UMAP neural network model is retained and a k-Nearest Neighbors (kNN) classifier is trained on the combination of marker expression and cell identities (see methods for details).

To illustrate the method, we used a dataset consisting of 10 PBMC samples from our previous work ^11^. A random set of nine PBMCs samples ^11^ was used to train the initial model and one independent sample was projected on the reference UMAP and annotation (**Figure 6c**). The projected data aligned well with the reference UMAP embedding as shown by a LISI score close to two demonstrating the desired mix between cells derived from the original embedding and the projected data (**Figure 6d**). Furthermore, the training of the kNN classifier resulted in an overall accuracy higher than 97% when predicting Phenograph clusters (**Figure S10a**) and 99% when predicting cell types (**Figure S11a**). The kNN classifier implementation in *cyCONDOR* also outputs the importance score calculated by the kNN model for each marker in the classification (**Figure S10b, S11b**) providing information on the relevance of each marker in the panel for the classification task. Label prediction based on the train classifier leads to a good overlap between the annotation of the training dataset and the new data (**Figure 6e, S9c**) When comparing the automated annotation provided by *cyCONDOR* with the manual annotation performed by annotating Phenograph clusters according to marker expression for the projected samples, we observe an almost perfect overlap (**Figure 6f**). Furthermore, also at the level of individual cell types and clusters a LISI score around two showed a good projection of the UMAP even for small clusters or minor cell types (**Figure S9d, S9e**). With this efficient approach, new samples can be automatically analyzed using a reference dataset without the need for manual annotation. As this process does not rely on the parallel processing of multiple samples, this analysis can be scaled indefinitely providing a robust framework for the analysis of thousands of samples and millions of cells over time even without the requirement of an HPC infrastructure. Considering the potential challenges in evaluating the expected variance in biological data, we envision our approach to be implemented incrementally. Initially, a reference dataset comprising a limited number of samples, designated as model V1, can be employed. While a small sample size may not fully encompass the entire range of human variation, as the number of samples increases, we anticipate developing an updated reference model, V2, to accommodate this expanded diversity. This incremental approach enables the continuous refinement of predictive accuracy.

### Harnessing machine learning for clinically relevant classification with cyCONDOR

Flow cytometry is commonly used as a clinical test for the diagnosis of several hematological diseases such as leukemia ^31^. Furthermore, in recent years, thanks to the advent of high dimensionality methodologies, HDC has been assigned great potential for the diagnosis of many other diseases (e.g. HIV, COVID-19, neurological diseases ^32^). Expanding from the use of a general model to project new samples (**Figure 6**), we implemented in *cyCONDOR* a set of functions to train clinical classifiers for the categorization of new samples without manual investigation (see methods for details - **Figure 7a, S12a**).

**Figure 7:**
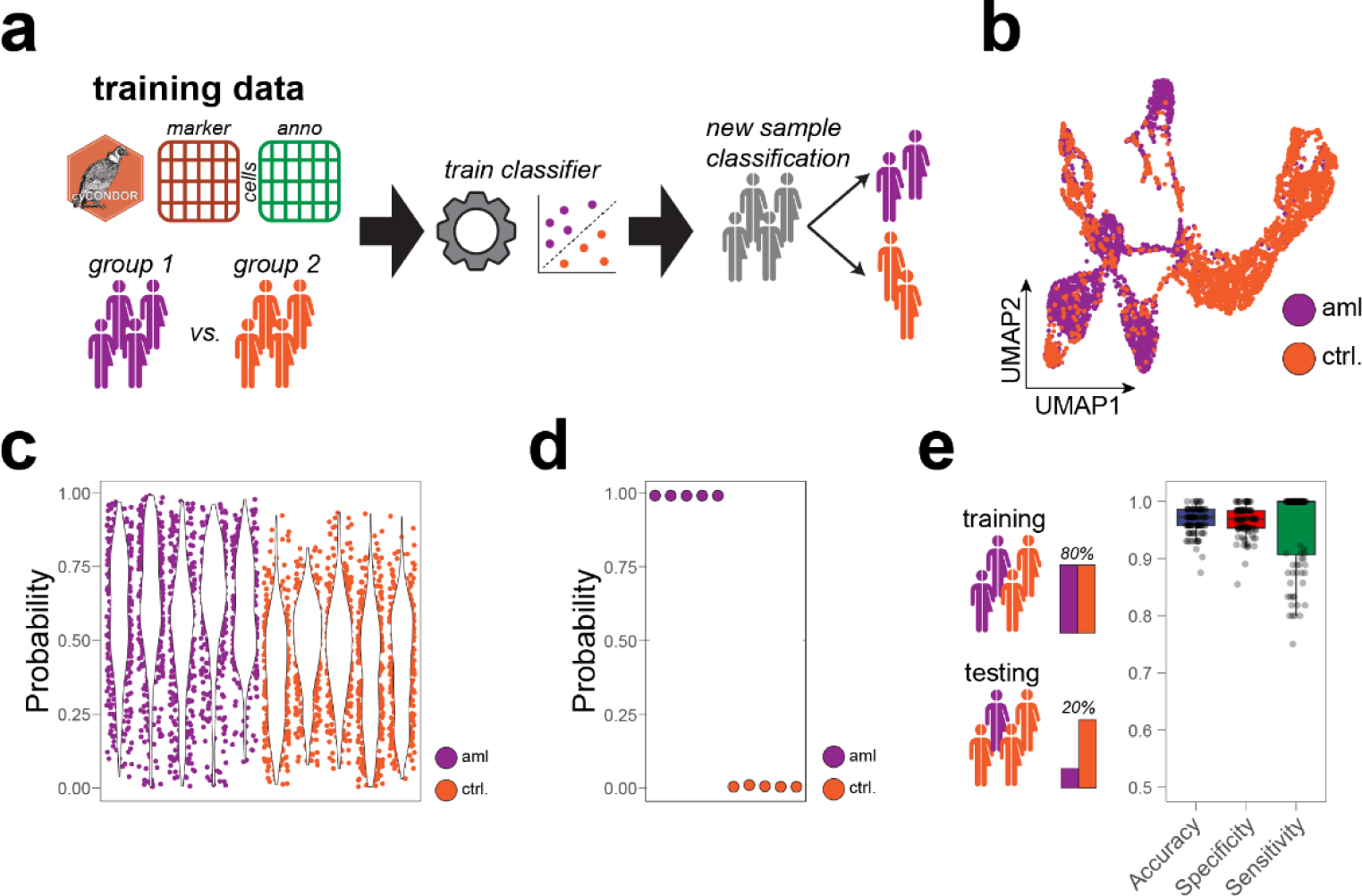
Direct implementation of clinical classifier allows the accurate classification of disease states. **a,** Schematic overview of the clinical classifier workflow implemented in *cyCONDOR*. **b,** UMAP visualization of the training dataset colored by experimental groups. **c,** Single-cell level probability for the test dataset split by sample and colored by experimental group. **d,** Sample level probability for the test dataset split by sample and colored by experimental group. **e,** Accuracy, specificity and sensitivity of a clinical classifier trained on the entire FlowCap-II dataset (100 permutations).

As a starting point for clinical classification tasks, we utilized the *CytoDx* model ^33^ which predicts clinical outcomes by individually assessing each cell’s association and averaging these signals across samples, and adapted it to the *cyCONDOR* ecosystem. To test the functionality of this module in *cyCONDOR*, we made use of the FlowCapII dataset, which serves as one of the gold-standard datasets for benchmarking machine learning (ML) classifiers on cytometry data ^34,35^. As a first step, we created a model using a selection of 20 samples from the *FlowCapII* dataset, which included samples from patients with acute myeloid leukemia (aml) and healthy control samples. We split the subset into a training dataset (5 *aml* and 5 *controls*) and a test dataset (5 *aml* and 5 *controls*). We first explored the difference between *control* and *aml* samples at the level of their UMAP embedding (**Figure 7b**) showing that cells from aml and control samples differentially populated the different subclusters. Independently from any cell type label, using a classification tree ^33^ we trained two classifiers, first at the level of individual cells (i.e. cellular classifiers **Figure S12b**), and consequently at the sample level (i.e. sample classifier **Figure S12c**). Already at the single-cell level, the classifier results showed a separation between *aml* samples and *controls* with an overall higher aml classification probability for aml-derived cells (**Figure S12b**). The aml model, derived by the decision tree algorithm was visualized as a tree map illustrating that the model can be visualized to allow further investigation of the decision-making processes employed by the classifier to assign a probability to each cell. As anticipated, the feature importance analysis for our cellular model showed markers of the myeloid lineage, such as *CD13*, as key determinants for classification (**Figure S12d**). For the sample classifier, the trained model was able to correctly classify the 10 samples used for training (**Figure S12c**). Next, the model was evaluated on the test dataset, which has no overlap with the training data, and we could see a similar increase in probability for aml-derived cells (**Figure 7c**) as well as a perfect classification of the 10 new samples at the sample level (**Figure 7d**). To extend the validation of the *cyCONDOR* framework for sample classification, we then included in the analysis the entire *FlowCapII* dataset, comprised of 359 samples (43 *aml* and 316 *controls*). We split this dataset into 80% training and 20% test data and randomized this selection 100 times to evaluate the real-world performances of our classifier (**Figure 7e**). Before training the training dataset of 80% of the data was balanced to have an equal number of *aml* and *control* cases while the test dataset was left unbalanced (*1 aml / 7.3 controls*) to reflect a real-world scenario. For each permutation, we calculated accuracy, specificity and sensitivity on the 20% test dataset showing optimal performance also on real-world data (**Figure 7e**). Collectively, *cyCONDOR* facilitates the classification of clinical HDC data on cellular and sample level, opening avenues for the widespread application of ML to HDC data.

## Discussion

Flow cytometry, developed in the early 1950s, has been a revolutionary technique for the understanding of heterogeneous tissues ^3^. It allows the quantification of multiple protein markers at single-cell resolution and can measure millions of cells in a single experiment ^3^. While recent advances in HDC have expanded the potential of cytometry to dissect complex tissues at the single-cell level ^36^, these advancements have also introduced a multitude of analytical challenges.

Traditional cytometry data analysis relies on the sequential selection of cells in two-dimensional plots (gating), which is adequate for a limited number of protein markers. However, as novel methodologies enable the simultaneous measurement of more than 50 proteins per cell, traditional analytical approaches become increasingly cumbersome and less effective.

In the last few years, several approaches besides commercial software have provided the cytometry community with tools to investigate HDC data using data-driven approaches commonly used by the single-cell transcriptomics community. *Cytofkit*, a pioneering project that ceased development in 2017, played a pivotal role in catalyzing a paradigm shift in the analysis of HDC ^7^. This tool has provided several data transformation and clustering approaches still used in the field ^7^. Other projects such as *SPECTRE* ^8^ and *Catalyst* ^9^ have increased the feature set available to the community by introducing approaches for signal overlap correction in CyTOF data ^37^ or computational pipelines for the analysis of CyTOF imaging datasets ^8^.

Complementary, several non-academic projects, such as *Cytobanks* or *Cytolytics* provide feature-rich tools, often with an intuitive graphical user interface (GUI) for the guided analysis of HDC data. Those tools are often able to handle small datasets with difficulties in scaling to larger ones, commonly produced with newer instruments. Accessibility to these pipelines is not free and necessitates access to external web servers, raising concerns about data privacy following national regulations ^38^.

In this study, we introduce *cyCONDOR* as an easy-to-use open-source ecosystem for HDC data analysis. Building upon existing tools like *SPECTRE*, *Catalyst* and *Cytofkit*, *cyCONDOR* prioritizes not only user-friendliness but also the biological interpretation of data with the scalability to millions of cells and the implementation of state-of-the-art ML methods. We first demonstrate the applicability of the *cyCONDOR* workflow to a broad range of data types including HDFC, CyTOF, Spectral Flow and CITE-seq (**Figure 2**). Furthermore, we showcase how *cyCONDOR* can efficiently mitigate the technical batch between datasets (**Figure 3**) and provide “publication-ready” comparisons between experimental groups (**Figure 5**). Most of these steps were already individually available in other analytical pipelines, nevertheless *cyCONDOR* focuses on the simplicity of use for non-computational biologist and offers better performance thanks to the implementation of multi-core computing for the most intensive steps (e.g. UMAP calculation or Phenograph clustering), drastically reducing computing times.

Additionally, *cyCONDOR* provides new analytical workflows aiming at the biological interpretation of the data and scalability to population-wide studies. In this manuscript, we demonstrate the application of *cyCONDOR* to investigate the continuous development of HSCs into the major immune cell lineages by inferring pseudotime (**Figure 4**). Moreover, the integration of a kNN classifier enables the projection of new data onto existing embeddings, facilitating limitless scalability of the *cyCONDOR* workflow and enabling continuous analysis of population-wide longitudinal studies (**Figure 6**). Furthermore, the possibility to easily train a clinical classifier within the *cyCONDOR* pipeline enables the applicability of *cyCONDOR* to clinical settings where sufficient data are available (**Figure 7**).

The focus of *cyCONDOR* on ease of use is still limited in some aspects. Cell type identification is still a laborious process and cannot be automated yet. When compared to single-cell transcriptomics where all transcripts are measured, HDC relies on a pre-selected set of markers. This pre-selection in the available parameter limits the use of reference mapping techniques such as *SingleR* and will still require manual annotation based on the marker expression. Future developments of *cyCONDOR* will provide the implementation of *Astir* ^39^, an interesting tool simplifying the process of cluster annotation.

Taken together, *cyCONDOR* provides an easy-to-use, end-to-end ecosystem for HDC data analysis extending on the available features of other approaches (**Figure S1**). We provide *cyCONDOR* as an open-source R package making it compatible with any common operating system (Mac OS, Windows and Linux). Furthermore, we provide *cyCONDOR* with a companion Docker Image ensuring full reproducibility of the analysis while costing only little computational overhead ^38^, simplifying the deployment of our tool, and limiting the risk of any incompatibility with other R packages.

## Methods

### Datasets

#### Chronic HIV, Human PBMCs, HDFC

The HDFC phenotyping data from control and chronic HIV donors ^11^ was kindly provided by Dr. Anna Aschenbrenner. Similarly to the SpectralFlow dataset reported above, debris were removed according to *FSC-A* and *SSC-A*, singlets were selected (*FSC-A vs. FSC-H*) and dead cells were removed. Compensated .fcs files were then exported for *cyCONDOR* analysis. This dataset was used to exemplify cyCONDOR capabilities for pre-processing (**Figure 2**), differential analysis (**Figure 5**) and data projection (**Figure 6**).

#### Rheumatoid Arthritis, Human whole blood, CyTOF

For the evaluation of the *cyCONDOR* ecosystem with CyTOF data (**Figure 2**), we downloaded the dataset reported by Leite Pereira et al. ^40^. From this dataset only healthy control 1 and 2 were used including both unstimulated and IL7 stimulated cells (*HEA1_NOSTIM.fcs, HEA1_STIM.fcs, HEA2_NOSTIM.fcs, HEA2_STIM.fcs*). The dataset was downloaded from FlowRepository (FR-FCM-Z293).

#### Healthy, Murine Spleen, SpectralFlow

For the evaluation of the *cyCONDOR* ecosystem with SpectralFlow data (**Figure 2**), we downloaded the dataset reported by Yang et al. ^41^. From this dataset we only used Spleen 1 and Spleen 2 (S1.fcs and S2.fcs). Before the analysis debris were removed according to *FSC-A* and *SSC-A*, singlets were selected (*FSC-A vs. FSC-H*) and dead cells were removed. Compensated .fcs files were then exported for *cyCONDOR* analysis. The dataset was downloaded from FlowRepository FR-FCM-Z4NB.

#### Healthy, Human PBMCs, CITE-seq

Healthy controls were collected as part of the DELCODE ^42^ study. PBMCs were stained with BD Rhapsody Ab-seq Immune Discovery Pannel kit (BD) according to manufacturer instructions. Raw sequencing reads were processed with the BD Rhapsody Pipeline (v.2.1) and UMI counts per cell were used for *cyCONDOR* analysis. Ab-seq counts were transformed with a Center log ratio transform (clr) before dimensionality reduction and clustering. This dataset was used to exemplify the use of *cyCONDOR* with CITE-seq data (**Figure 2**).

#### Healthy, Human PBMCs, HDFC

Healthy controls were collected as part of the DELCODE ^42^ study and measured over several days with a BD Symphony S6 cell sorter. Similarly to the SpectralFlow dataset reported above, debris were removed according to *FSC-A* and *SSC-A*, singlets were selected (*FSC-A vs. FSC-H*) and dead cells were removed. Compensated .fcs files were then exported for *cyCONDOR* analysis. This dataset was used to exemplify the batch correction workflow implemented in *cyCONDOR* (**Figure 3**).

#### Healthy, Bone Marrow, CyTOF

The CyTOF dataset reported by Benadll and colleagues ^26^ was downloaded from CytoBank. Before cyCONDOR analysis the data was cleaned as described in the CytoBank analysis. Shortly singlets were selected according to cell length and 191-DNA staining. The surface staining for bone marrow 1 was used for the analysis (*Marrow1_00_SurfaceOnly.fcs*). With this dataset we exemplify the trajectory inference and pseudotime capabilities of *cyCONDOR* (**Figure 4**)

##### AML, FC - FlowCap-II

The FlowCap-II AML dataset ^34,35^ was downloaded from FlowRepository (FR-FCM-ZZYA). For the evaluation of the performances of cyCONDOR clinical classifier all samples from panel 4 were used without any further processing. We use this dataset to benchmark the machine learning classifier implemented in *cyCONDOR* (**Figure 7**).

### Structure of the *cyCONDOR* object

We developed the *cyCONDOR* ecosystem as an R package. The current version of the *cyCONDOR* package (v 0.1.5) was developed with R v 4.3.0 and Bioconductor v 3.17. The *cyCONDOR* object, containing all the data resulting from a *cyCONDOR* analysis is structured as an R list with separate data slots for marker expression (*expr*), cell annotation (*anno*), dimensionality reduction (*pca*, *umap*, *tsne*), and clustering (*clustering*). Individual elements are structured as R data frames with each row representing an individual cell and each column a parameter. The structural integrity of the *cyCONDOR* object can be evaluated at each step with built-in functions to ensure the object was correctly manipulated.

### Data pre-processing and transformation

Individual *.fcs* files are imported in R and merged with the sample annotation using the *prep_fcd* function. This function imports each *.fcs* or *.csv* files, merges all expression tables into a single data frame and performs an autologicle transformation ^7,43,44^ marker-wise. Before merging, each cell is assigned a unique cell name composed of the name of the file or origin and sequential numbering. Additionally, a cell annotation table is initialized from a provided sample metadata table. The output *cyCONDOR* object will contain both data frames, the transformed expression data frame and the annotation data frame, and will be used for all the downstream processes.

### Dimensionality reduction

*cyCONDOR* provides several functions to perform different types of dimensionality reductions, each function requires a *cyCONDOR* object and outputs a *cyCONDOR* object including the coordinates of the reduced dimension for each cell. Except for the PCA, all other dimensionality reductions provided with *cyCONDOR* (UMAP, tSNE and DM) can use as input the principal components (recommended option shown in this manuscript) or the marker expression. The user can also decide the number of PCs to use for the calculation to reduce the computational requirements.

#### Pseudobulk principal component analysis (PCA)

To calculate the pseudobulk principal components the *runPCA_pseudobulk cyCONDOR* function calculates at first the mean marker expression across all cells. The resulting matrix is then used to perform a PCA. As the dimensionality of the output matrix differs from the dimensionality of the *cyCONDOR* object, only in this case the output of the function will not be the modified *cyCONDOR* object but a new list comprising only the PCA coordinates and the input dataset.

#### Principal Component Analysis (PCA)

The *cyCONDOR runPCA* function uses the *prcomp* base R function to compute the principal components for each cell. The output of the function is the original *cyCONDOR* object extended by the PC coordinates.

#### Uniform Manifold Approximation and Projection (UMAP)

The *cyCONDOR runUMAP* function uses the *uwot* UMAP implementation (CRAN). Compared to other R native implementations of the UMAP algorithms this implementation allows parallelizing the UMAP calculation, enabling high performances and allows to retain the neural network model, which is used to project new data to existing UMAP embeddings (see section “*Data projection*” below). The output of the function is the original *cyCONDOR* object extended by with the UMAP coordinates.

#### t-Distributed Stochastic Neighbor Embedding (tSNE)

The *cyCONDOR* function *runtSNE* relies on the *Rtsne* implementation of the tSNE algorithm to calculate this non-linear dimensionality reduction. Similarly to the UMAP calculation, the output is the original *cyCONDOR* object added with the tSNE coordinates.

#### Diffusion Map (DM)

To calculate a diffusion map, the *cyCONDOR* function *runDM* relies on the *destiny* package^45^. Similar to the other dimensionality reduction approach this function will output the original *cyCONDOR* object extended by the DM coordinates.

### Clustering

#### Phenograph

Phenograph clustering is performed with the *Rphenoannoy* R package which compared to the original R implementation ^45^ allows parallelization of Phenograph calculation. For applying the *cyCONDOR* function *runPhenograph* the user will provide a *cyCONDOR* object and decide which data to use for Phenograph clustering (usually PCA). The function will return a *cyCONDOR* object including the result of the clustering algorithm. The user can also optimize the *k* parameter to generate a more broad or fine-grained clustering.

#### FlowSOM

FlowSOM clustering is performed with the *FlowSOM* R package ^16^. With the *cyCONDOR* function *runFlowSOM* the user will provide a *cyCONDOR* object and decide which data to use for FlowSOM clustering (usually PCA). The function will return a *cyCONDOR* object including the results of the clustering algorithm. The user also needs to provide the number of final clusters as input.

#### Batch correction

The *cyCONDOR* ecosystem implements *harmony* ^19^ to account for differences between experimental batches. The implementation of harmony provides the option to correct experimental batches at both the levels of marker expression with the function *harmonize_intensities* and principal components with the function *harmonize_PCA*. The output of both options can be used to calculate a non-linear dimensionality reduction and clustering. While this is technically possible it is not advisable to use the harmonized marker expression for differential expression analysis as this might lead to overestimation or underreppresentation of the differences. For both functions, the output will consist of the original *cyCONDOR* object with the addition of the harmonized valuse.

#### Pseudotime analysis

*cyCONDOR* implements *slingshot* ^25^ for pseudotime analysis and trajectory inference. After data pre-processing including transformation, dimensionality reduction, clustering and cell annotation, the function *runPseudotime* takes the coordinates of a dimensionality reduction (e.g. PCA or UMAP) to infer pseudotime and trajectories. The *runPseudotime* function also requires a vector with the cell labels. Within the *runPseudotime* function the user can define fixed starting and ending points for the trajectory. Additionally, *cyCONDOR* offers a user-friendly validation option that recalculates the trajectory using each cluster/metacluster as the starting point. This functionality aids in identifying the best-fitting model for any given cell differentiation task. Pseudotime and trajectories can be easily visualized with *cyCONDOR* built in functions.

#### Data projection

The workflow for the projection of new data to an existing reference dataset consists of two main steps. First, the preparation of the reference dataset consists of the training of the UMAP neural network and retaining the model within the *cyCONDOR* object with the *runUMAP* function setting *ret_model* to *TRUE*. After annotation of the dataset, a kNN classifier is also trained on the reference data using as input the expression values and the cell labels of each cell. This step is performed with the *cyCONDOR* function *train_transfer_model* which takes advantage of the *caret* framework for machine learning in R ^46^. The kNN model will also be retained within the *cyCONDOR* object. For the projection of new data, the functions *learnUMAP* and *predict_labels* will take the built models from the reference dataset to project the new cells into the existing UMAP embedding and to predict the cell labels.

#### Clinical classifier

With the *cyCONDOR* implementation of the *CytoDx* ^33^ model it is possible to easily train a machine-learning (ML) classifier. The *cyCONDOR* function *train_classifier_model* takes as input a *cyCONDOR* object (expression values) and a variable defining the different categories to train a classifier of both individual cells and samples. The performance of the classifier can be easily exploited with the pre-build function as well as the decision tree used for the classification ^33^. The output of this function will be the original *cyCONDOR* object with the addition of the ML model.

For the classification a of new samples, the *predict_classifier* function takes as input the *cyCONDOR* object containing the samples to classify and the pre-trained model (stored in the training *condor* object). The output of this function will be the *cyCONDOR* object added with the probability of the classification for each cell and each sample.

### Statistical analysis and data visualization

Statistical significance was calculated in R (v. 4.3.0) with an unpaired two-sided t-test if not stated differently. A p-value < 0.05 was considered significant. All data were visualized using R (v. 4.3.0) with the packages ggplot2, pheatmap or the built-in functions of *cyCONDOR* (v. 0.1.4). cyCONDOR implements several statistical testing methods for the comparison between groups, the function *boxplot_and_stats*, can calculate a *t-test* or *wilcox-test* when comparing two groups or in case of more then two groups and *anova* or *kruskal* test, for *t-test* or *wilcox-test* the user can define if the sample are paired. All box plots were constructed in the style of Tukey, showing median, 25^th^ and 75^th^ percentiles; whisker extends from the hinge to the largest or lowest value no further than 1.5 ∗ IQR from the hinge (where IQR is the interquartile range, or distance between the first and third quartiles); outlier values are depicted individually. Confusion matrices were used to show relative proportion across groups as a fraction of samples from the respective condition contributing to each cluster or cell type.

## Declarations

### Availability of data and materials

All data used in this manuscript are publicly available as described in the individual figure. R environment necessary to reproduce the analysis shown in this manuscript will be made available upon publication on Zenodo (LINK, DOI).

### Code availability

*cyCONDOR* source code is available on GitHub (https://github.com/lorenzobonaguro/cyCONDOR). All code to reproduce the analysis shown in this manuscript is available on GitHub (https://github.com/lorenzobonaguro/cyCONDOR_reproducibility). The data reported in this manuscript were analyzed with *cyCONDOR* v0.1.4 and Bioconductor 3.17.

## Authors’ contributions

Conceptualization was by L.B, T.P., M.B and J.L.S. The methodology was devised by L.B, S.M, C.K, J.L., T.K., C.C. and L.B. S.M., C.K., J.L., T.K., S. W., T.Z. performed formal analysis. J.L., T.K., T.Z. carried out the investigations. The draft manuscript was written by L.B, M.B. and J.L.S. All authors reviewed and edited the manuscript. Visualization was by L.B. S.M. and C.K.. The project was supervised by L.B. and T.P.. Funding acquisition was by L.B, T.P., M.B and J.L.S.

---

**Figure S1.**
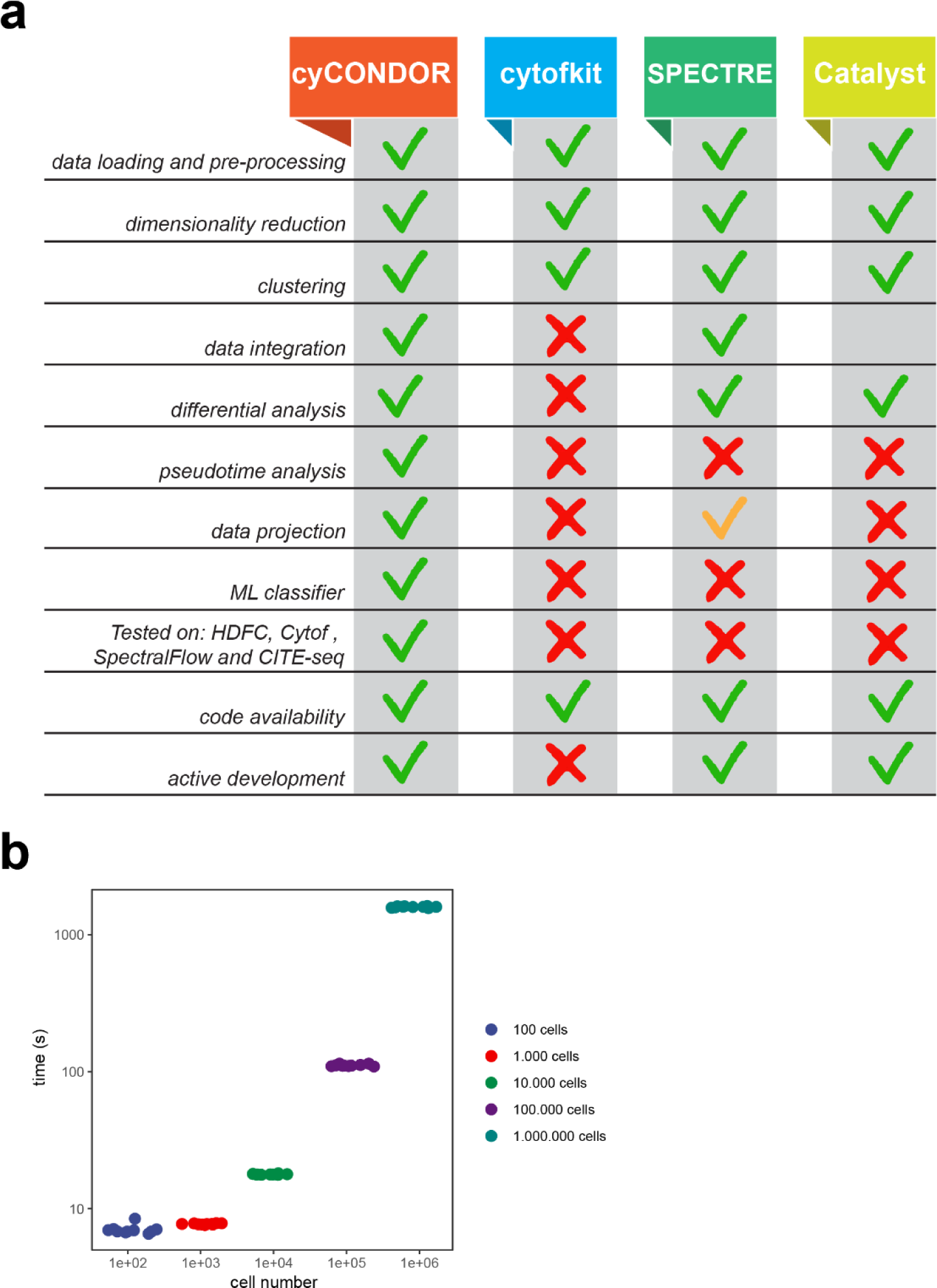
**a,** Comparative table of *cyCONDOR* with the most diffused cytometry data analysis frameworks. **b,** Performance analysis of the cyCONDOR workflow; data loading and transformation, PCA, UMAP and Phenograph clustering were performed on different numbers of cells (100, 1000, 10000, 100000, 1000000) for ten times each. The result show linear scaling of the cyCONDOR ecosystem. All measurement were taken using the cyCONDOR Docker image on a Windows 10 workstation equipped with Intel Core i7-8700K CPU and 32 Gb or system memory.

**Figure S2.**
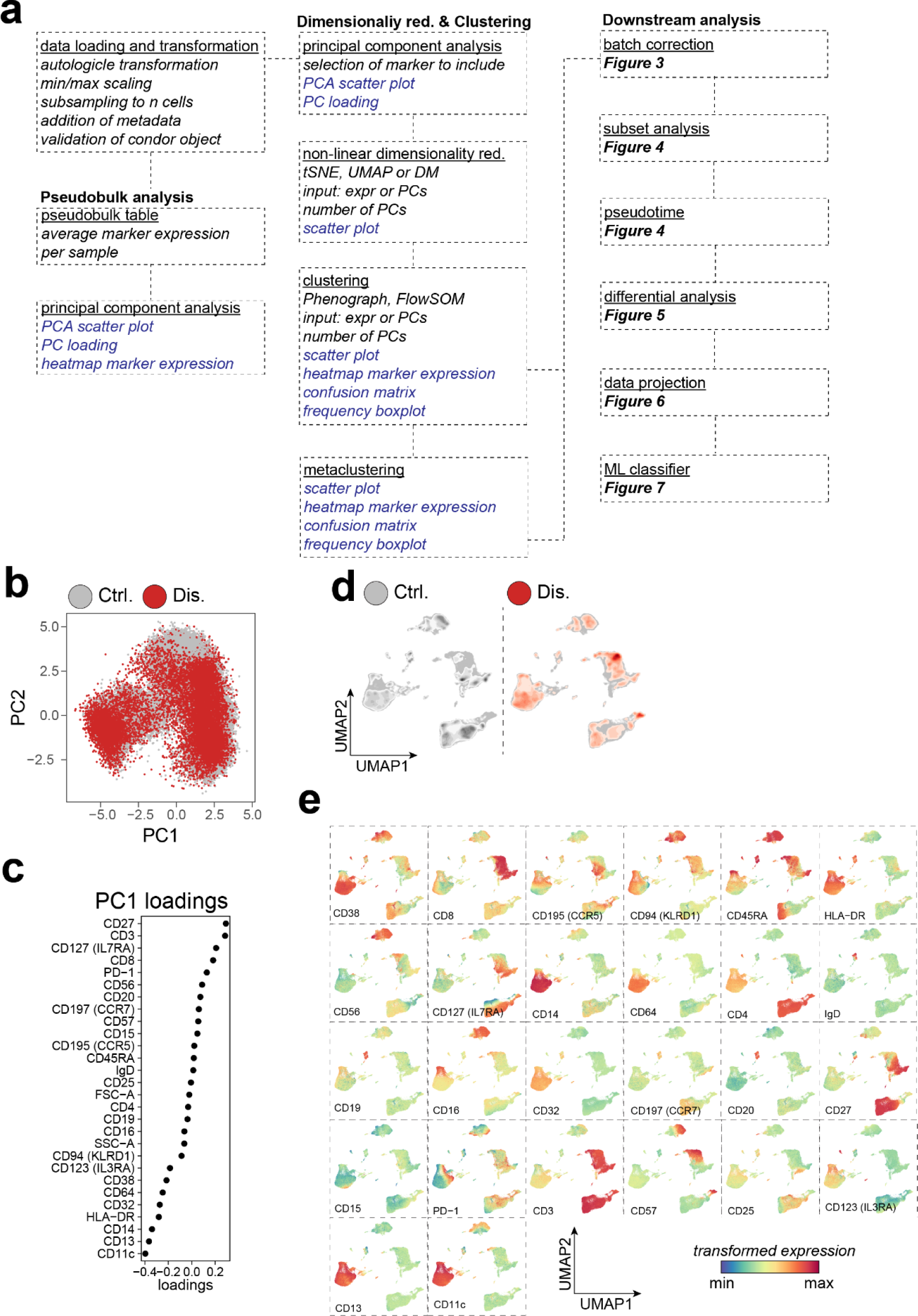
**a,** Detailed schematic of *cyCONDOR* preprocessing and downstream analysis. **b,** Scatterplot of single-cell level PC coordinates colored by experimental group. **c,** Loading of the first PC. **d,** UMAP colored according to the density of cells of each experimental group. **e,** UMAP colored according to the transformed expression of each marker in the dataset.

**Figure S3.**
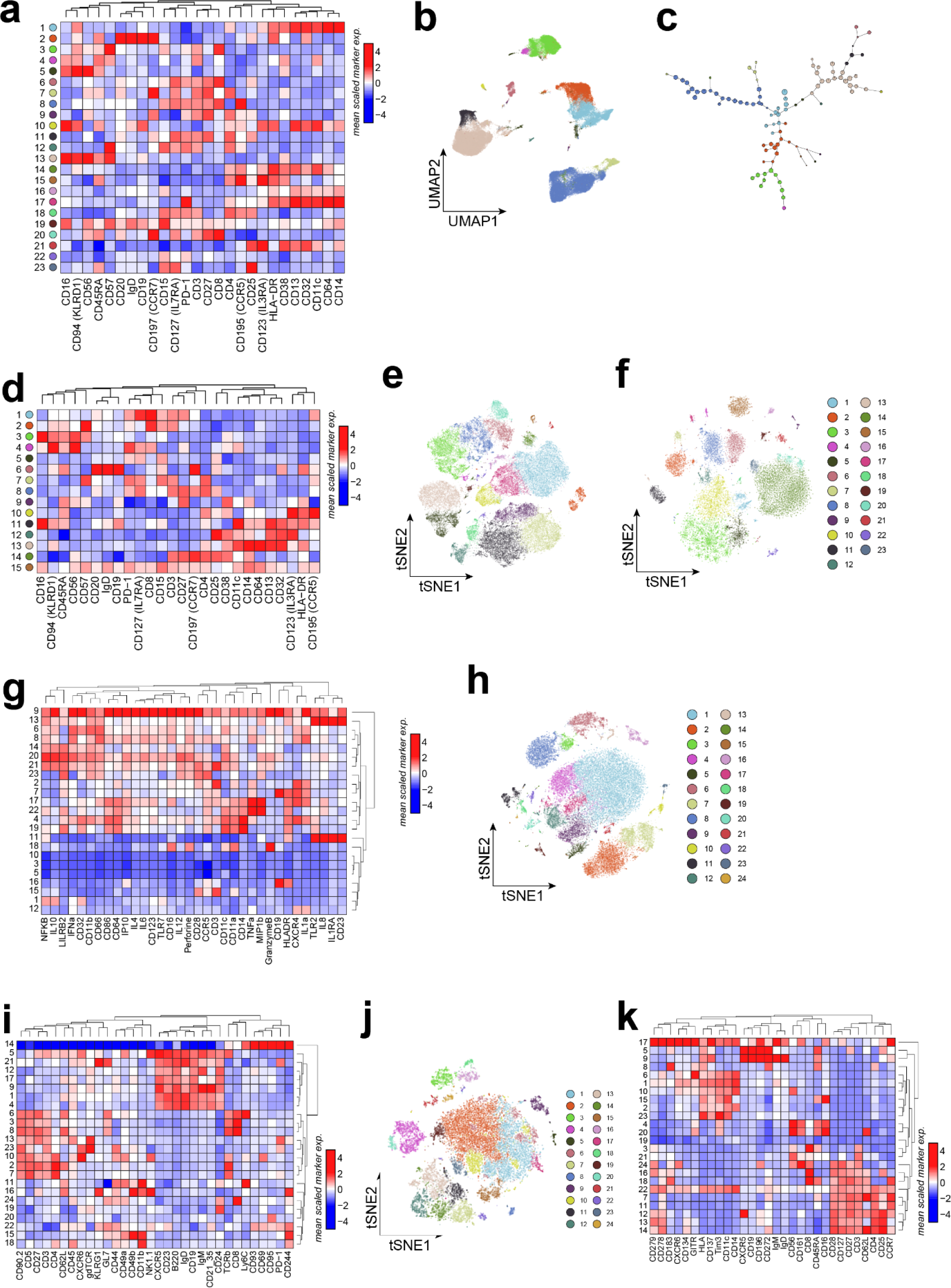
**a-e,** visualization of HDFC data (chronic HIV dataset). **a,** Heatmap of the average expression of each marker split by Phenograph cluster. b, UMAP colored according to FlowSOM clustering. **c,** SOM visualization colored by FlowSOM clustering. **d,** Heatmap of the average expression of each marker split by FlowSOM cluster. **e,** tSNE visualization of HDFC data colored by Phenograph clustering. **f,** tSNE visualization of CyTOF data colored by Phenograph clustering. **g,** Heatmap of the average expression of each marker split by Phenograph cluster, CyTOF data. **h,** tSNE visualization of SpectralFlow data colored by Phenograph clustering. **i,** Heatmap of the average expression of each marker split by Phenograph cluster, SpectralFlow data. **j,** tSNE visualization of CITE-seq data colored by Phenograph clustering. **k,** Heatmap of the average expression of each marker split by Phenograph cluster, CITE-seq data.

**Figure S4.**
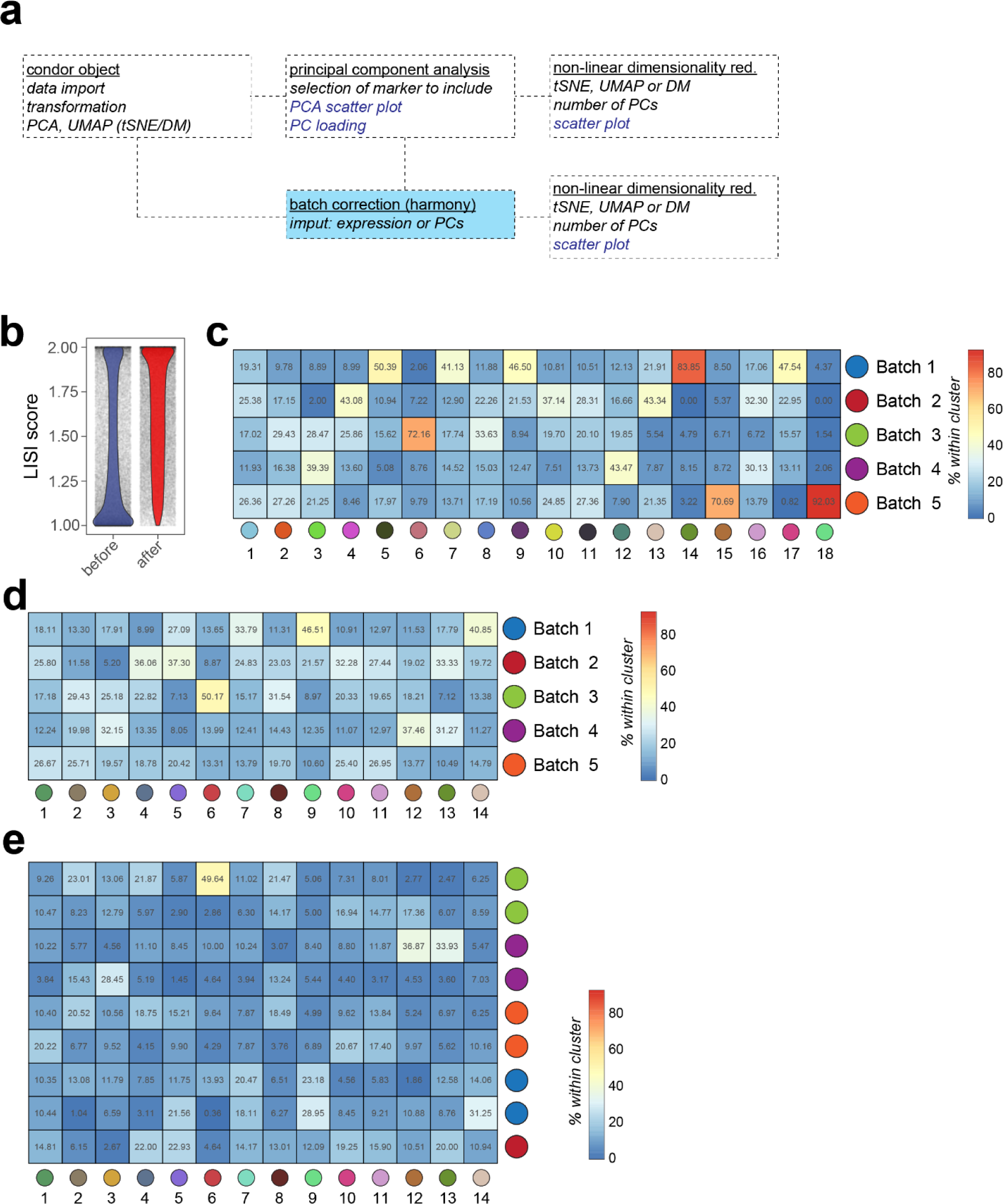
**a,** Detailed schematic of the batch correction workflow implemented in *cyCONDOR*. **b,** LISI score between batches before and after batch correction**. c,** Confusion matrix of the Phenograph clusters (not corrected data) split by experimental batch. **d,** Confusion matrix of the Phenograph clusters (corrected data) split by experimental batch. **e,** Confusion matrix of the Phenograph clusters (corrected data) split by sample.

**Figure S5.**
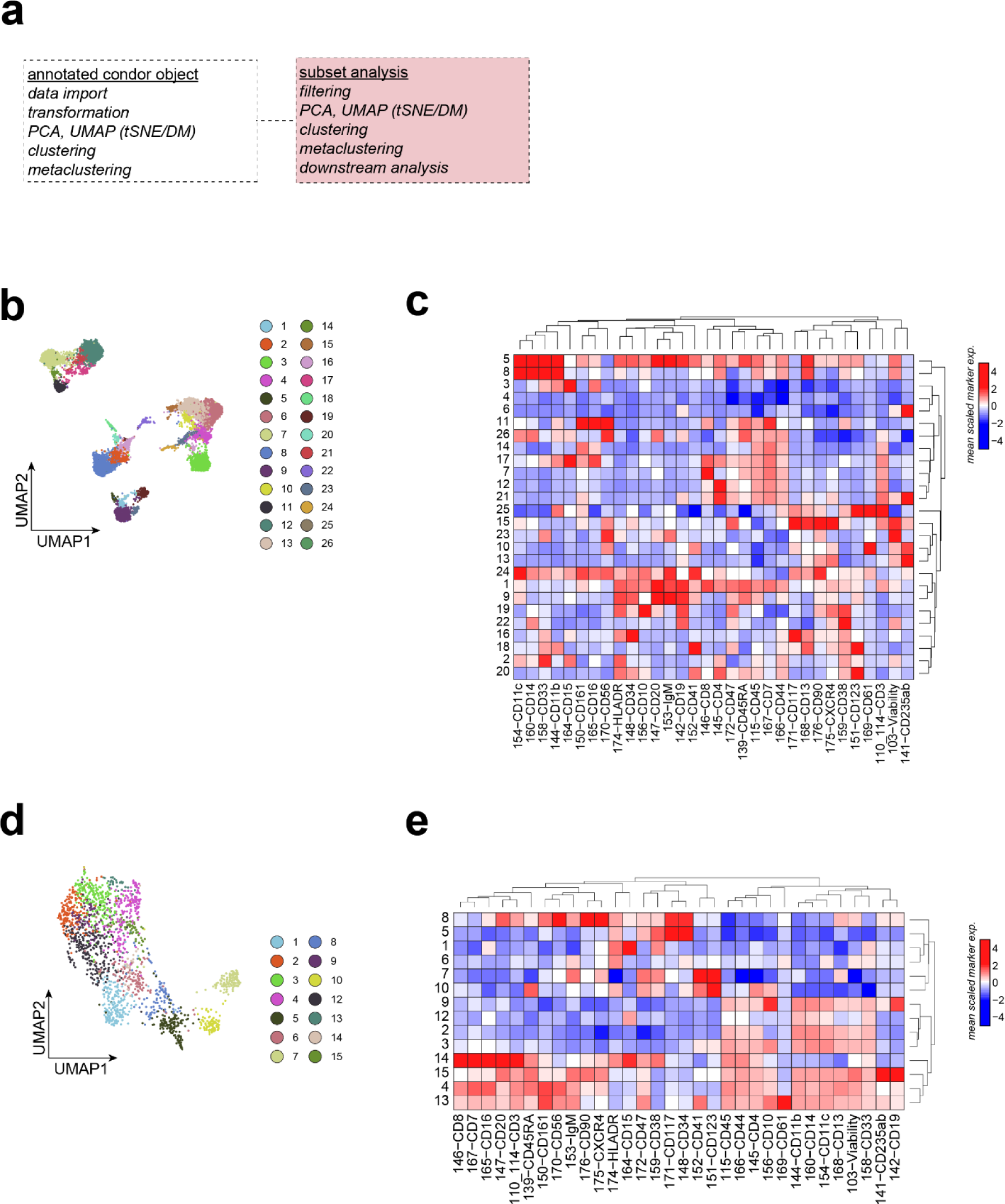
**a,** Detailed schematic of the subsetting workflow implemented in *cyCONDOR*. **b,** UMAP of all bone marrow cells colored according to the assigned Phenograph cluster. **c,** Heatmap of the average expression of each marker split by Phenograph cluster. **d,** UMAP of the subsetted dataset colored according to the newly assigned Phenograph cluster. **e,** Heatmap of the average expression of each marker split by Phenograph cluster, subsetted dataset.

**Figure S6.**
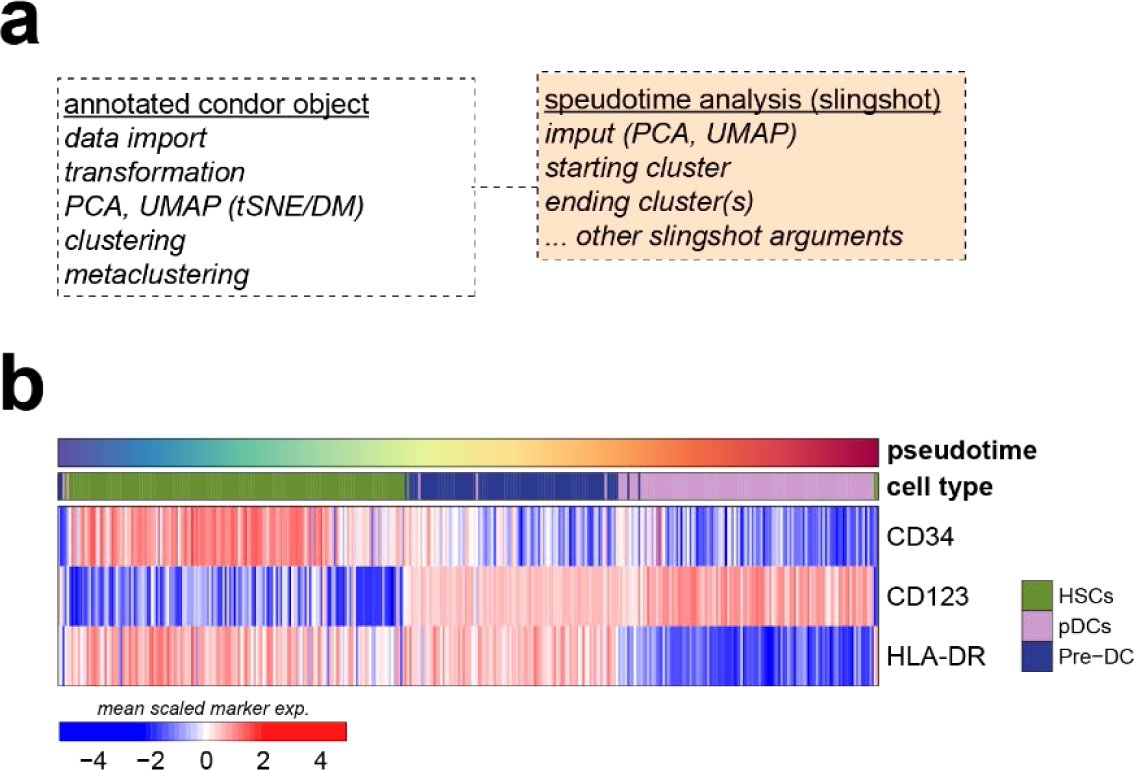
**a,** Detailed schematic of the pseudotime inference workflow implemented in *cyCONDOR*. **b,** Heatmap marker expression in cells belonging to the pDC trajectory ordered according to the inferred pseudotime.

**Figure S7.**
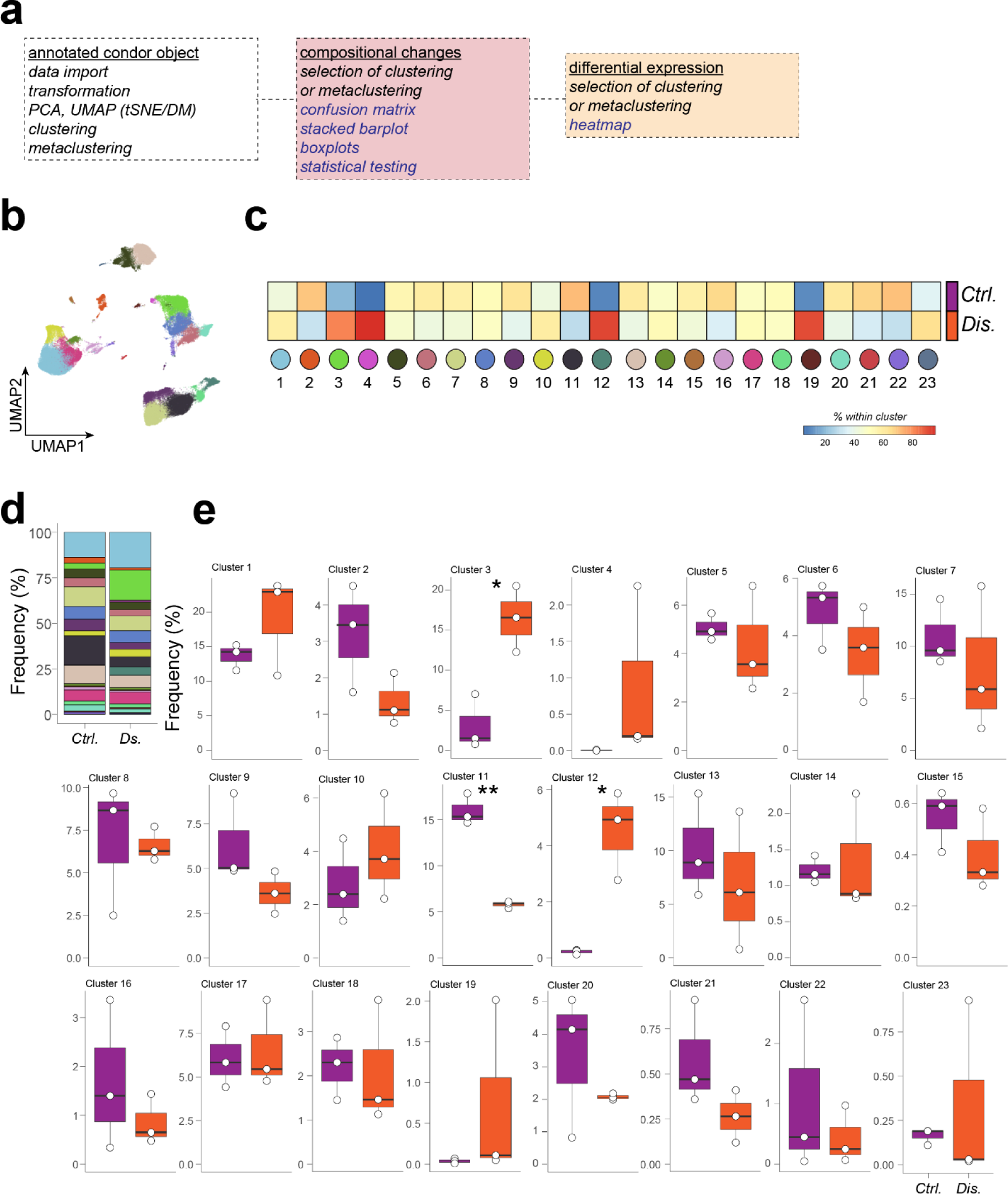
**a,** Detailed schematic of the differential analysis implemented in the *cyCONDOR* ecosystem. **b,** UMAP colored by Phenograph clustering. **c,** Confusion matrix of the Phenograph clusters split by experimental group. **d,** Stacked barplot of the Phenograph clusters frequencies split by experimental groups. **e,** Boxplot of the frequency of each Phonograph cluster split by experimental group.

**Figure S8.**
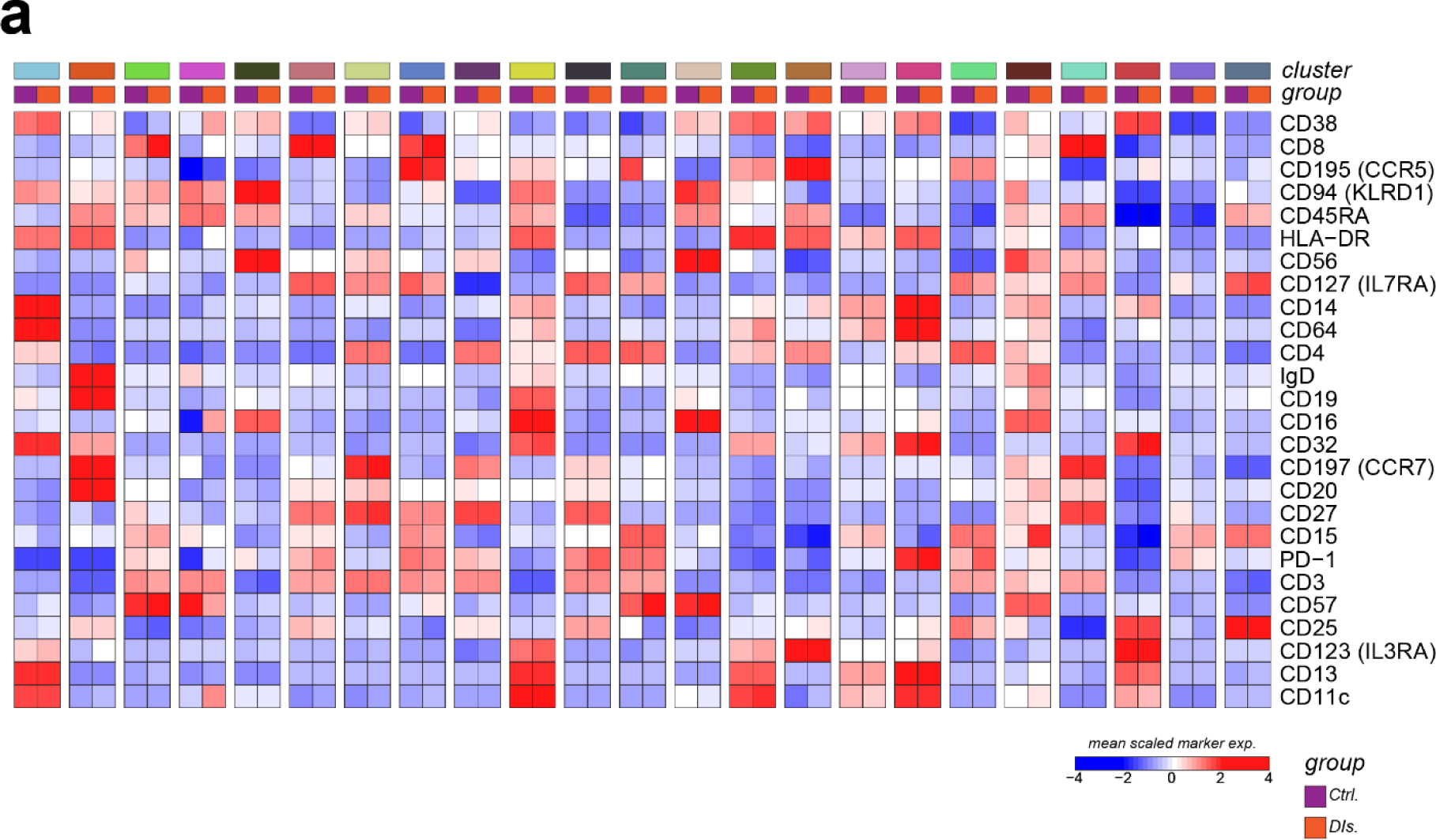
**a,** Heatmap of the average expression of each marker split by Phenograph cluster and experimental group.

**Figure S9.**
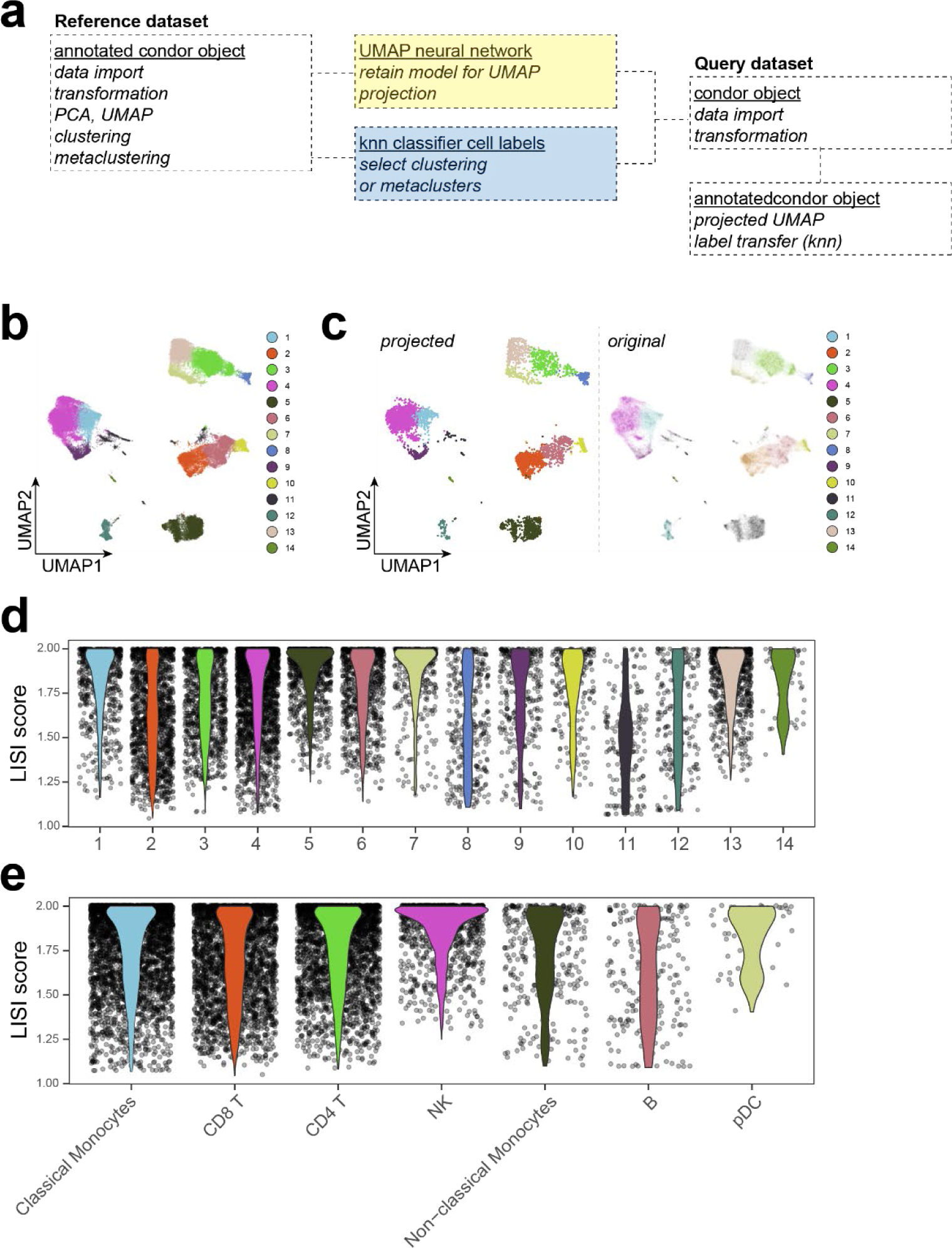
**a,** Detailed schematic of the data projection workflow implemented in *cyCONDOR*. **b,** UMAP visualization of the training dataset colored according to the assigned Phenograph clustering. **c,** UMAP visualization of the projected data colored according to the predicted Phenograph cluster. **d,** LISI score calculated between training data and projected data split by Phenograph cluster. **e,** LISI score split by assigned cell type.

**Figure S10.**
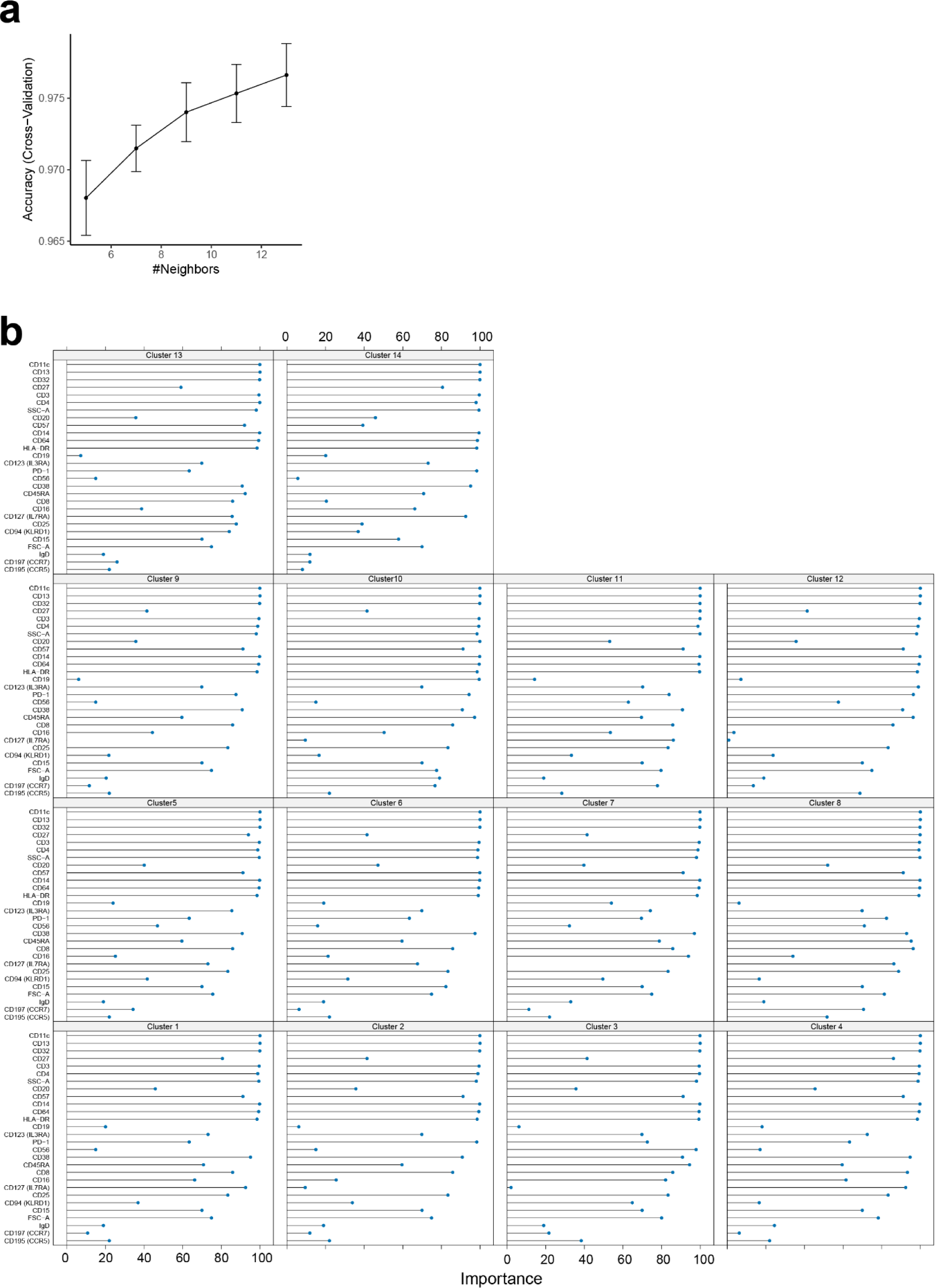
**a,** Accuracy of the kNN model trained to predict Phenograph clusters across different numbers of neighbors used for model optimization. **b,** Importance score for the assignment of each cell label for individual Phenograph clusters.

**Figure S11.**
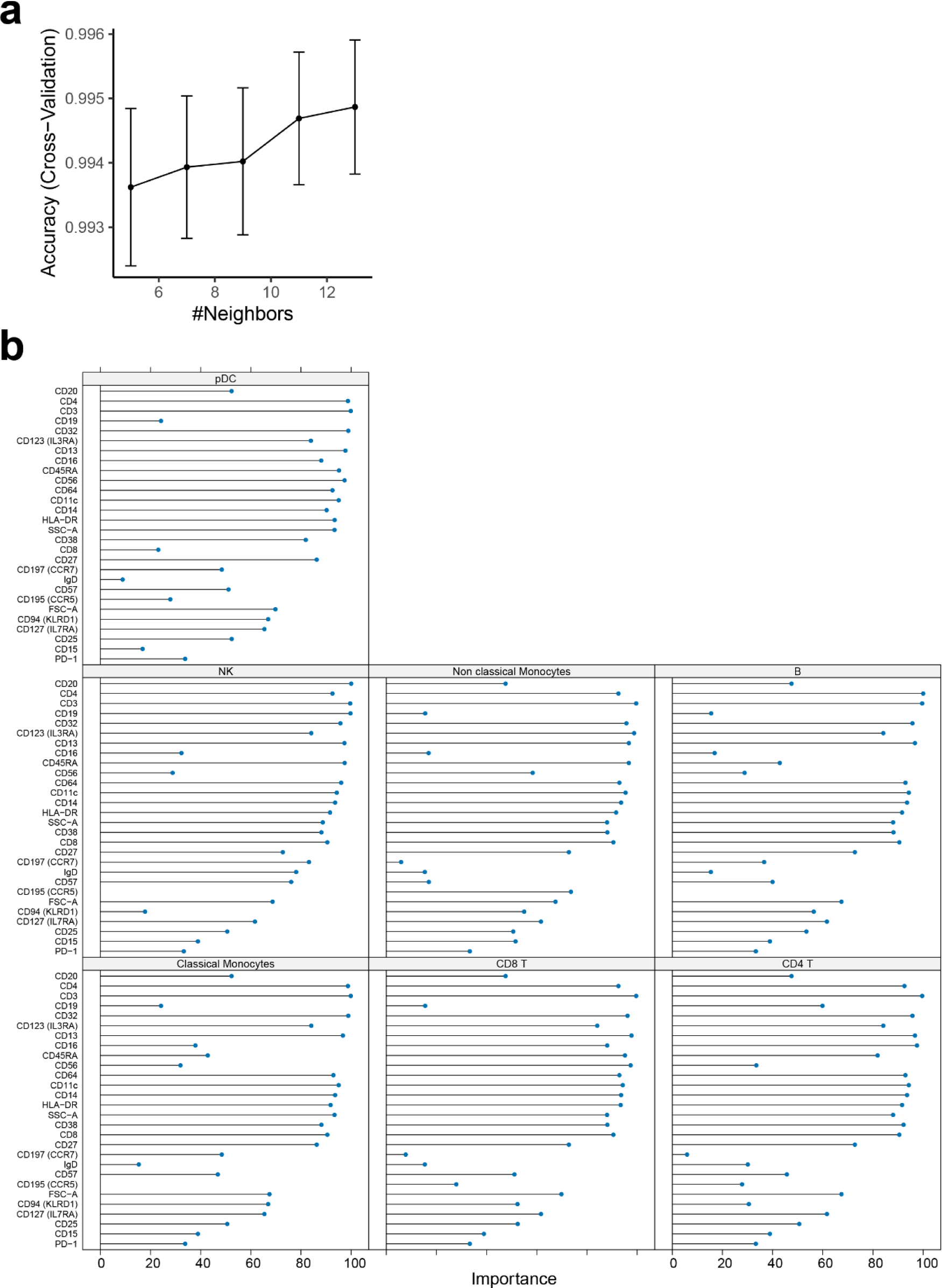
**a,** Accuracy of the kNN model trained to predict annotated cell labels across different numbers of neighbors used for model optimization. **b,** Importance score for the assignment of each cell label for individual annotated cell labels.

**Figure S12.**
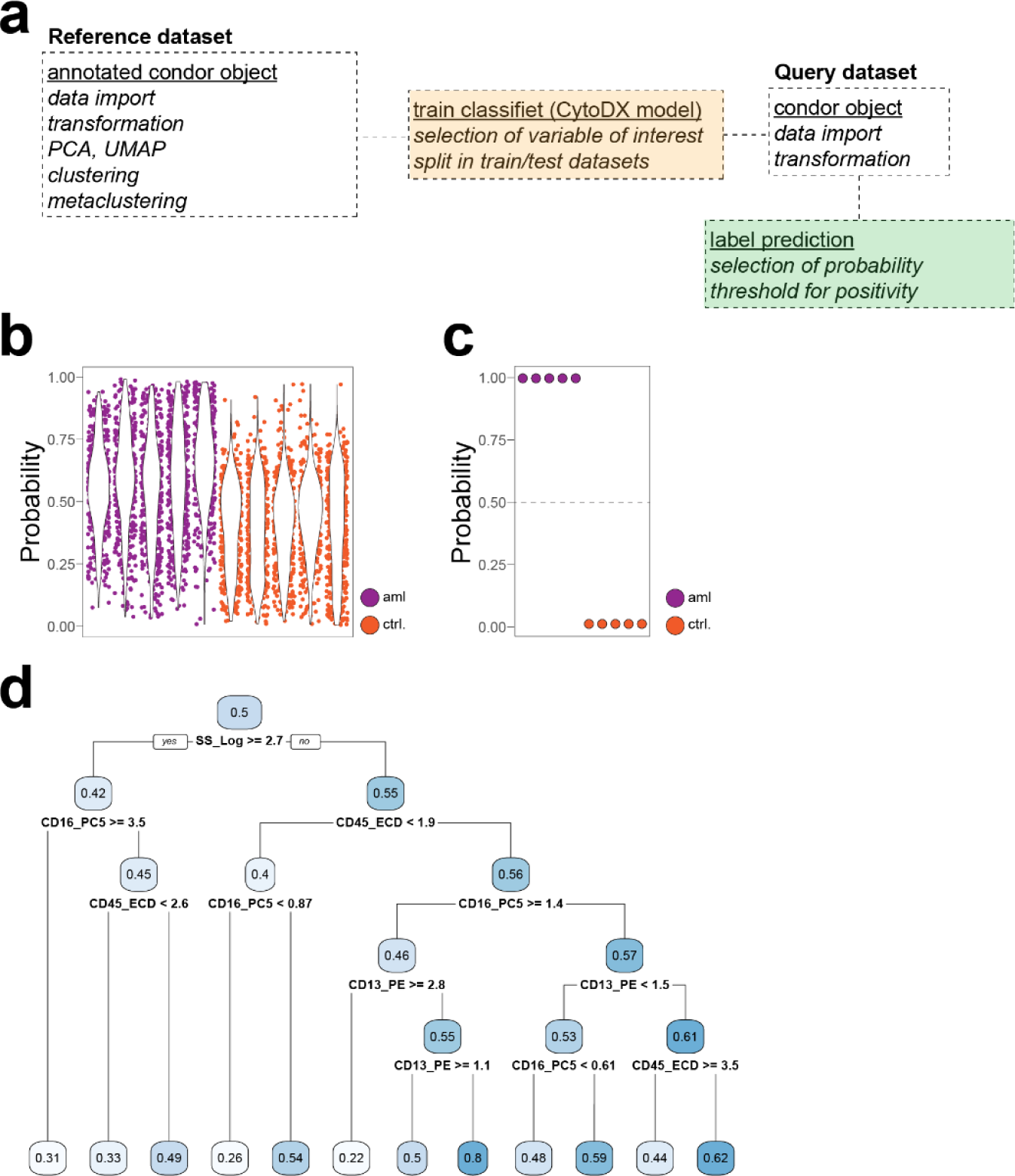
**a,** Detailed schematic of the clinical classifier workflow implemented in *cyCONDOR*. **b,** Single-cell level probability for the training dataset split by sample and colored by experimental group. **c,** Sample level probability for the training dataset split by sample and colored by experimental group. **d,** A decision tree classifies cells as aml. Each branch of the tree represents a specific characteristic, and the value at each node shows the likelihood of aml association for that group of cells. The rules at each branch further divide the cell population into more refined subgroups based on additional characteristics.

